# Biomarkers indicate altered outer retinal fluid homeostasis in the absence of Kv2.1/Kv8.2 channels

**DOI:** 10.64898/2026.06.27.734996

**Authors:** Joseph G. Laird, James Soetedjo, Shivangi M. Inamdar, Alex R. Bock, Ehimenmen S. Ataman, Miles A. Pufall, Bruce A. Berkowitz, Sheila A. Baker

## Abstract

**Purpose:** Photoreceptor Kv2.1/Kv8.2 voltage-gated potassium channels carry an outward potassium current, helping to set the resting membrane potential and to shape dim light responses. Because potassium flux in the outer retina influences extracellular osmolarity and fluid distribution, we hypothesized that Kv2.1/Kv8.2 channels also contribute to fluid homeostasis in this region of the retina.

**Methods:** OCT imaging was performed in Kv8.2 heterozygous (Het) and knockout (KO) mice aged ∼5 weeks under dark- and light-adapted conditions. Light-dark differences in the distance between the external limiting membrane (ELM) and retinal pigment epithelium (RPE) (ΔELM-RPE) were calculated to quantify light-evoked outer retinal expansion, a biomarker associated with changes in subretinal space (SRS) fluid handling. As a secondary outcome, outer nuclear layer (ONL) thickness was also measured under both lighting conditions. Retinal gene expression was assessed by RNA-seq and droplet digital RT-PCR. Retinal protein expression was determined by western blotting and immunolabeling.

**Results:** ΔELM-RPE was significantly reduced in Kv8.2 KO mice compared with Het controls, indicating altered light-evoked outer retinal dynamics compatible with changes in subretinal space fluid handling. ONL thickness exhibited a small but significant light-dark change that was different between genotypes. Transcriptomic analyses revealed upregulation of osmosensitive genes, including osmolyte transporters and aquaporins. AQP1 protein expression in photoreceptors increased.

**Conclusions:** These findings are consistent with altered outer retinal fluid homeostasis following loss of Kv8.2 and support a model in which photoreceptor potassium efflux contributes to osmotic water movement into the subretinal space.

## Introduction

Kv2 voltage-gated potassium channels are widely expressed and conduct delayed rectifier currents in many excitable cells. In photoreceptors, Kv2.1 assembles with the regulatory subunit Kv8.2 to form heterotetrameric channels that carry the potassium current *I*_Kx_.^1,2^ *I*_Kx_ constitutes the major component of the outward arm of the circulating dark current, contributing to the establishment of the resting membrane potential and shaping dim light responses. Loss of Kv8.2 disrupts *I*_Kx_ causing cone dystrophy with supernormal rod responses (CDSRR), also known as KCNV2 retinopathy.^3^ This inherited retinal disease is characterized by delayed and reduced ERG responses, early loss of central cone vision with macular atrophy, and variable loss of night vision. In mice, loss of Kv8.2 causes delayed and reduced ERG responses characteristic of CDSRR, and rod photoreceptors undergo a slow degeneration.^4–7^ It is not known if the photoreceptor degeneration associated with loss of Kv8.2 is a direct result of disrupted photosignaling or if there are additional physiological consequences to altered potassium homeostasis in the outer retina.

Another role for potassium flux in the outer retina besides regulating photoreceptor membrane voltage is to regulate osmotic fluid uptake from the subretinal space (SRS). The subretinal space constitutes a restricted extracellular microenvironment that is the interface between photoreceptors and the RPE. There is substantial net flux of fluid from the neural retina, through the SRS, and across the RPE into the choroidal circulation as this is the route for the removal of excess water produced by the prodigious metabolic rate of photoreceptors.^8–15^ The process of fluid absorption from the SRS by the apical RPE is coupled to the concentration of potassium in the SRS through the activity of the Na⁺- K⁺- Cl⁻ cotransporter (NKCC1).^16,17^ Unlike mechanisms governing water clearance from the SRS, the processes that facilitate movement of water from photoreceptors into the SRS are less well studied.

Kv2.1/Kv8.2 channels are positioned along the photoreceptor inner segment plasma membrane so that they carry a flow of potassium from photoreceptors into the SRS.^1,7^ Therefore, it is possible that potassium flux through these channels contributes to drawing excess water out of photoreceptors into the SRS. This idea predicts that in Kv8.2 knockout (KO) mice, photoreceptors would retain water, and the SRS would become relatively dehydrated.

Measuring photoreceptor volume directly is not feasible. However, changes in SRS fluid dynamics can be assessed noninvasively using an OCT imaging-based biomarker, “ELM-RPE.” OCT resolves the ELM and RPE as distinct outer retinal bands, allowing measurement of the distance between them (ELM-RPE thickness) and providing a measure of light-evoked outer retinal dynamics that previous studies have linked to changes in SRS hydration. This interval includes the photoreceptor inner and outer segments and the extracellular SRS so includes both outer retinal cellular anatomy and SRS volume. Importantly, ELM-RPE thickness changes reproducibly with extended light adaptation: it contracts in the dark and expands in the light.

Prior microelectrode and multimodal imaging experiments showed that the light-evoked expansion corresponds with changes in the volume of the SRS and outer retinal water content and mobility.^18–26^ The light-evoked expansion of the ELM-RPE reported by OCT develops over minutes to hours and is distinct from the rapid and transient outer segment swelling thought to occur in response to phototransduction-induced osmotic changes that are measured by optoretinography.^20,26^

The physiological basis of the light-evoked expansion of ELM-RPE thickness has been linked to photoreceptor metabolism and pH-sensitive fluid transport by the RPE.^27^ In darkness, increased photoreceptor metabolism generates CO [through mitochondrial respiration and lactate through aerobic glycolysis. The resulting acidification of the SRS stimulates fluid removal by the RPE.^8^ In light, reduced photoreceptor metabolism diminishes this stimulus, slowing fluid removal such that the SRS volume ─ and hence ELM-RPE thickness ─ expands.^18–26^

In this study, we quantified paired light-dark changes in ELM-RPE thickness in Kv8.2 KO mice using OCT and assessed osmosensitive responses at the transcript and protein levels. We found that Kv8.2 KO mice exhibit reduced light-evoked ELM-RPE expansion, upregulation of osmosensitive genes, and increased AQP1 protein expression. Together, these findings support a model in which Kv2.1/Kv8.2-mediated potassium flux participates in maintaining fluid homeostasis within the outer retinal microenvironment.

## Materials and Methods

### Animals

Kv8.2 KO (*Kcnv2*^-/-^) mice (MGI:7660814) were maintained in the laboratory. Mice were housed in a central vivarium, maintained on a standard 12/12-hour light/dark cycle, with food and water provided ad libitum in accordance with the Guide for the Care and Use of Laboratory Animals of the National Institutes of Health. Routine genotyping was performed by Transnetyx. The gene-expression experiments described below were performed before the breeding strategy was modified to reduce animal numbers and therefore used WT controls; subsequent imaging and protein experiments used Het littermates. WT and Het mice were indistinguishable in ERG responses and longitudinal ONL thickness (Supplemental Figs. S1–S2), supporting their interchangeable use as controls. A mix of male and female mice were used in all experiments. For experiments requiring anesthesia, a mixture of ketamine (87.5 mg/kg) and xylazine (2.5 mg/kg) was used with body temperature maintained by keeping animals on a heating pad. Eye drops to dilate the pupils (1% tropicamide) and maintain hydration (Genteal) were applied. All procedures were approved by the University of Iowa IACUC committee and adhered to the ARVO Statement for the Use of Animals in Ophthalmic and Vision Research.

### Optical Coherence Tomography (OCT)

For light-adapted imaging, mice were exposed to room lighting (∼800 lux) for one hour while awake and then anesthetized immediately before imaging. For dark-adapted imaging, mice were dark-adapted overnight and anesthetized under very dim red light (<1 lux). Mice rested for one week between sessions and were then imaged under the opposite lighting condition. Both sessions occurred between P33 and P41, with an average imaging age of 5 weeks. A total of 6 Kv8.2 Het and 11 Kv8.2 KO animals (representing 3 different litters) were used. Images were collected with a Bioptigen spectral-domain imaging system (model #75-10023 B00, Bioptigen, Inc.) as previously described.^7^ The central 100 B scans for each image file were then sent to Wayne State University for image analysis.

A first-pass rigid body registration with RNiftyReg (a function in R) was used to rotate the image and interpolate the signal at each pixel. Next, non-rotational rigid body approaches (at the level of a given row or column of pixels) were applied three times. The 100 images were then visually compared as a final step before averaging. After registration, layer segmentation boundaries were estimated using a machine learning model–based computer program, as described previously.^27^ Boundaries were first estimated with a previously described U-net convolutional neural network trained using the “dice loss” function and the Adam optimizer (learning rate = 0.001), with 665 previously labeled images for training and 166 and 356 images for validation and testing, respectively. Model predictions were then postprocessed by applying a shortest-path algorithm. Based on these model-based estimates for laminar borders, final segmentation refinement was performed with an R script, as described previously.^27^ Once segmented, inferior and superior retinas (350–624 µm from the optic nerve head on the inferior and superior sides) were each analyzed; starting at 350 µm ensured that our data were analyzed away from the optic nerve head, where the outer retina is relatively uniform in all OCT data. We measured the ONL and ELM-RPE thicknesses from in-house R scripts that objectively extracted layer boundaries obtained after searching the space provided by machine-learning estimates (“seed boundaries”) as above. The ELM and RPE are initially identified by local signal maxima and the R script determines the ELM-RPE thickness by calculating the distance from ELM to the basal side of the RPE at the level of Bruch’s membrane.^28,29^ Complete statistical outputs for the OCT analyses, including model summaries, estimated marginal means, and contrasts (Tables S1A–S4C), are provided in Supplemental File S1.

### RNA-seq

Mice were euthanized by CO₂ inhalation followed by cervical dislocation. After enucleation and removal of the anterior segment, neural retinas were dissected in PBS free of the RPE, choroid, and sclera. The two retinas from each mouse were pooled to generate one biological sample. Retinas were collected from three Kv8.2 WT and five Kv8.2 KO mice at 1 month of age. Total RNA was isolated from retinas collected into RNAlater using the RNeasy Mini Kit (Qiagen) according to the manufacturer’s protocol. Library preparation and sequencing were performed by Novogene.

Sequencing reads were adapter-trimmed using TrimGalore and quantified against the mouse transcriptome (GENCODE Mus musculus GRCm38) using Salmon (v1.10.2) with sequence-, GC-, and positional-bias correction. Transcript-level abundances were summarized to gene-level counts using tximport. Before normalization and modeling, expression of *Kcnv2* was examined to confirm the knockout in samples designated as KO. One sample (KO2) showed no reduction in *Kcnv2* expression relative to the WT range and was excluded from all downstream analyses. The final analysis therefore included 3 WT and 4 KO biological samples.

Differential gene expression analysis was performed in R using DESeq2. Low-count genes were filtered, and one surrogate variable estimated using surrogate variable analysis was included as a covariate to account for expression variation unrelated to genotype. The model tested the effect of genotype while adjusting for this surrogate variable. Log₂ fold change estimates were shrunk using apeglm, and genes with a Benjamini–Hochberg-adjusted p < 0.01 were considered differentially expressed. Complete RNA-seq quality-control results, bioinformatic workflow, analysis parameters, and supporting analyses are provided in Supplemental File S2. RNA-seq data have been deposited in the Gene Expression Omnibus (GEO) under accession number GSE327286.

### RT-ddPCR

Separate RNA preparations from Kv8.2 WT and KO mice at 3.5 months of age were used for reverse transcriptase (RT) droplet digital PCR (RT-ddPCR), with cDNA pooled within genotype and assayed at four input amounts (1, 5, 10, and 20 ng), using probe sets for mouse *Slc6a6* (Mm.PT.58.11922589) and *Nfat5* (Mm.PT.58.7500558) purchased from IDT, as previously described.^30^

### Western blotting

Neural retinas were collected as described above; two retinas from each mouse were pooled per sample. Retinas from 3 Kv8.2 Het and 3 Kv8.2 KO mice at 2 months of age were collected. Retinas were homogenized in 300 µl lysis buffer (1% CHAPS, PBS, protease inhibitor cocktail), clarified by centrifugation at 5,000 × g for 5 minutes and the supernatant mixed with 4x LDS sample buffer and 10x DTT, without heat denaturation. Samples (20 µl) were separated on 4-20% TGX gels and transferred to PVDF for blotting. REVERT 700 Total Protein Stain was used for lane normalization followed by incubation in Licor blocker. Primary antibodies were anti-AQP1 (Abcam, Cat# ab168387, RRID:AB_2810992) used at 1:4000 and anti-AQP4 (Millipore Cat# AB3594, RRID:AB_91530) used at 1:5000; secondary antibody was anti-rabbit: DyLight800 (Rockland, Cat# 5151P) used at 1:15000. Blots were imaged using a LI-COR Odyssey and band intensity analyzed with Image Studio (v5) software.

### Immunolabeling

Posterior eyecups from 3 Kv8.2 Het and 3 Kv8.2 KO mice, at 2 months of age, were dissected in PBS, fixed in 4% paraformaldehyde (prepared in PBS), cryoprotected in 30% sucrose, and embedded in tissue freezing media. Sets of Kv8.2 Het and KO retina sections were collected onto the same slides to ensure they were treated identically. After blocking with 10% normal goat serum containing 0.5% triton X-100, sections were incubated overnight at 4°C with anti-AQP1 (Abcam, Cat# ab168387, RRID:AB_2810992) used at 1:2000. After rinsing with PBS, sections were incubated with secondary antibody (anti-rabbit IgG: Alexa 647 (ThermoFisher, Cat #A-21235, RRID:AB_2535804) used at 1:500, mixed with Hoechst 33342, for two hours at room temperature. Images were collected using a THUNDER Imager 3D Tissue Fully automated upright research microscope Leica DM6 B equipped with a Leica DFC9000 GT camera. Image analysis, including THUNDER computational clearing and adjusting intensity levels for display, was performed using LASx software. Image acquisition and display settings were held constant for each Kv8.2 Het-KO pair. Signal intensity in the inner segment region was measured using FIJI. Nonspecific blood vessel-associated signal and signal from presumed AQP1-positive red blood cells were outside the region of interest and were not included.

### Mechanical isolation of rod outer segments (OS)

Posterior eyecups from 4 Kv8.2 Het and 4 Kv8.2 KO mice (from 2 litters), at 5-6 weeks of age, were dissected in mouse Ringers (130 mM NaCl, 3.6 mM KCl, 2.4 mM MgCl_2_, 1.2 mM CaCl_2_, and 10 mM HEPES, pH 7.4). Individual retina were gently shaken for 15 sec in 300 ul of 4% paraformaldehyde (prepared in mouse Ringers) containing 2 µg/ml WGA-Alexa 594 (ThermoFisher, catalogue number W11262), onto a mixed cellulose ester membrane then incubated 20 minutes at room temperature. Fluid was drawn off and replaced with a drop of Fluormount-G before adding a coverslip. Images were acquired at 40x using the THUNDER microscope described above. 50 intact OS were measured per animal; intact was defined by the presence of a sphere of retained inner segment material at one end with a long, unbroken rod-shaped extension.^31^

### Statistical Analysis

OCT biomarker data, ELM-RPE and ONL were analyzed using a linear mixed model with fixed effects of genotype, condition, side, and all interactions, and a random intercept for each mouse nested within genotype (biomarker ∼ genotype x condition x side + (1| genotype:mouse)). For ΔELM-RPE and ΔONL, the lighting condition is already encoded in the outcome so a mixed model with fixed effects of genotype, side, and their interaction (biomarker ∼ genotype x side + (1 | genotype:mouse)) was used. For all models, denominator degrees of freedom were estimated using the Satterthwaite approximation. Estimated marginal means and pairwise contrasts were computed and visualized in R (v4.4.1), with p values adjusted for multiple comparisons using the Holm method; results were visualized using Quarto (v1.5.45). For RT-ddPCR, normalized copy number was modeled separately for each transcript by linear regression with cDNA input, genotype, and their interaction as fixed effects. Differences between the slopes of the pooled WT and KO dilution series were evaluated using a t-test of the cDNA input × genotype interaction coefficient. Western blot band intensities, immunolabeling signal intensities,and isolated rod OS lengths were quantified from independent biological samples (with technical replicates averaged) and compared between genotypes using unpaired two-sample t-tests with Welch’s correction. Generative AI (ChatGPT-5, OpenAI) was used to assist with R scripting.

### Data and Code Availability

OCT image-processing and segmentation code is available at https://github.com/hm5798/Kv_Paper_Sheila. OCT statistical analysis code, and the OCT measurement data used in those analyses along with RNA-seq analysis code are available at https://github.com/sheilainiowa/Kv82_biomarkers.

## Results

### Reduced light-evoked ELM-RPE expansion in Kv8.2 KO mice

We used OCT to measure the distance between the ELM and RPE bands (ELM-RPE thickness), as well as ONL thickness, in the inferior and superior retina of mice approximately five weeks of age (Fig. 1A). Representative paired OCT images were obtained from the inferior retina of Kv8.2 Het and KO mice after one hour of light adaptation and after overnight dark adaptation (Fig. 1B). From these images it is apparent that for the Het mice, the ELM-RPE is thicker in the light than the dark as expected, while for the KO mice there is a much smaller difference in ELM-RPE due to lighting conditions. Quantification of ELM-RPE thickness was calculated separately for the inferior and superior retina for a group of 6 Het and 11 KO animals (Fig. 1C). In the inferior retina, ELM-RPE was thicker in the light adapted images relative to the dark adapted images by 3.65 µm in Het mice (95% CI [2.62, 4.68], p < 0.001) but only by 0.90 µm in Kv8.2 KO mice (95% CI [0.14, 1.67], p = 0.021). ELM-RPE thickness was also lower in Kv8.2 KO mice than in Het controls under both light-adapted (mean difference 6.23 µm, 95% CI [4.95, 7.51]) and dark-adapted conditions (mean difference 3.48 µm, 95% CI [2.20, 4.76]; both p < 0.001). The same overall pattern was observed in the superior retina: light adaptation increased ELM-RPE thickness in both genotypes, but the increase was smaller in Kv8.2 KO mice (Fig. 1C; Tables S1D–S1E).

**Figure 1.**
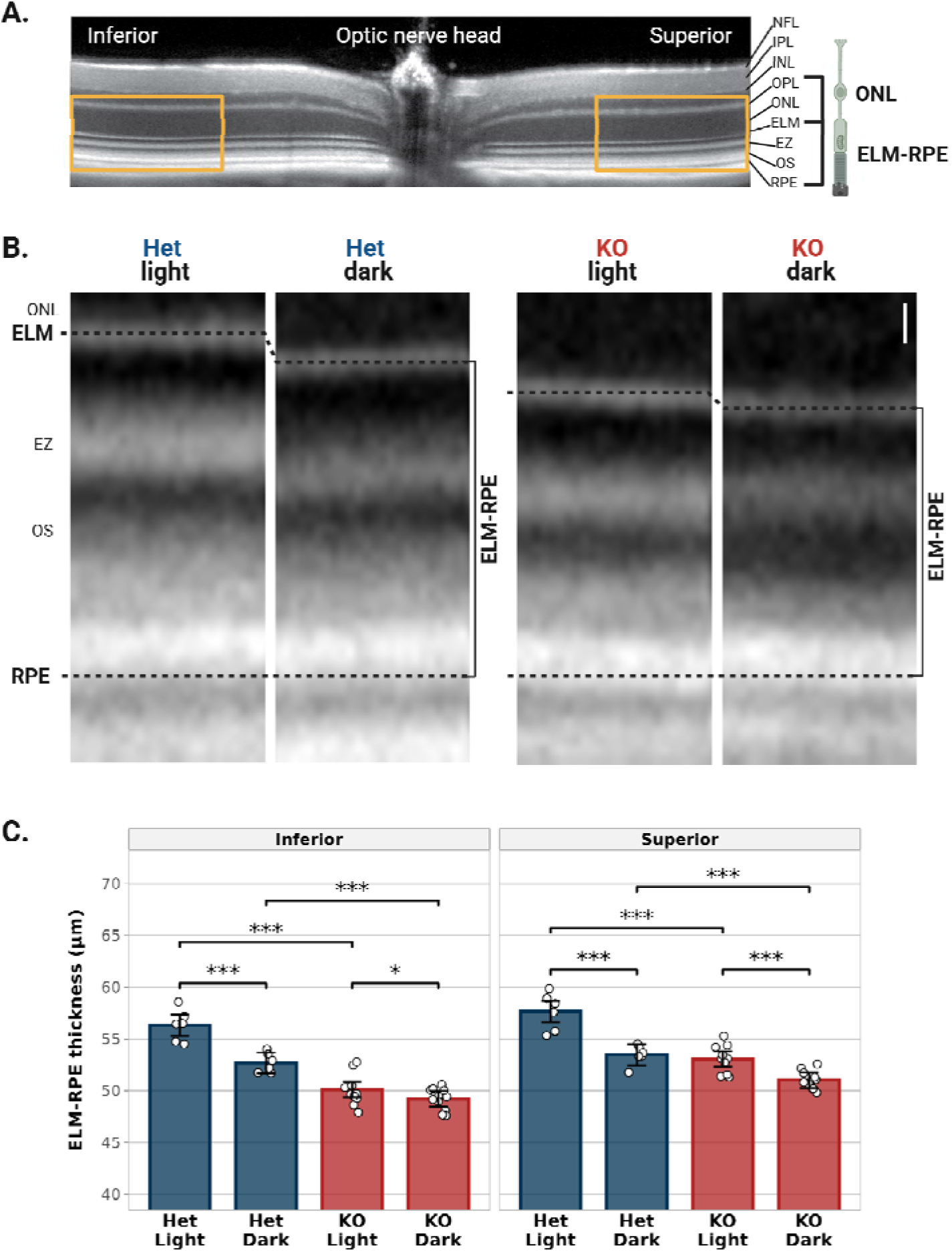
Altered light-dependent ELM-RPE thickness in Kv8.2 KO retina. (**A**) Representative OCT image showing inferior and superior retinal regions analyzed relative to the optic nerve head (orange rectangles). Retinal layers are labeled, and brackets indicate the anatomical boundaries of the ONL and ELM-RPE measurements. (**B**) Representative paired light- and dark-adapted OCT images from individual Kv8.2 Het and Kv8.2 KO mice. Scale bar, 12 µm. Dashed lines indicate the ELM and RPE boundaries used to measure ELM-RPE thickness, and brackets indicate the measured distance. All images are from the inferior retina. (**C**) Quantification of ELM-RPE thickness in the inferior and superior retina under light- and dark-adapted conditions for each genotype. Points represent individual mice; bars show estimated marginal means ± 95% CI (Kv8.2 Het, blue, n = 6; KO, red, n = 11). Brackets indicate pairwise comparisons (*p < 0.05, ***p < 0.001). Abbreviations: NFL, nerve fiber layer; IPL, inner plexiform layer; INL, inner nuclear layer; OPL, outer plexiform layer; ONL, outer nuclear layer; ELM, external limiting membrane; EZ, ellipsoid zone; OS, outer segments; RPE, retinal pigment epithelium.

We considered the possibility that structural changes could impact ELM-RPE thickness. Therefore, we next examined ONL thickness in the same OCT images (Fig. 2A). ONL thickness in these images was reduced in Kv8.2 KO mice compared with Hets (12% in inferior retina and 8% in superior retina in the light-adapted state), indicating retinal degeneration had begun with the amount of thinning consistent with only the loss of approximately one row of photoreceptor nuclei (Fig. 2A, Table S3C-S3D). To test if the extent of retinal degeneration at this age was accompanied by shortening of outer segments, which would impact our interpretation of ELM-RPE thickness, we mechanically isolated rod OS to measure their length independent of the inner segment and SRS. Isolated outer segments tended to be slightly shorter in Kv8.2 KO mice than in Het controls, but the difference was not statistically significant (mean difference 1.23 µm, 95% CI [-0.61, 3.08], p = 0.125; Fig. 2B).

**Figure 2.**
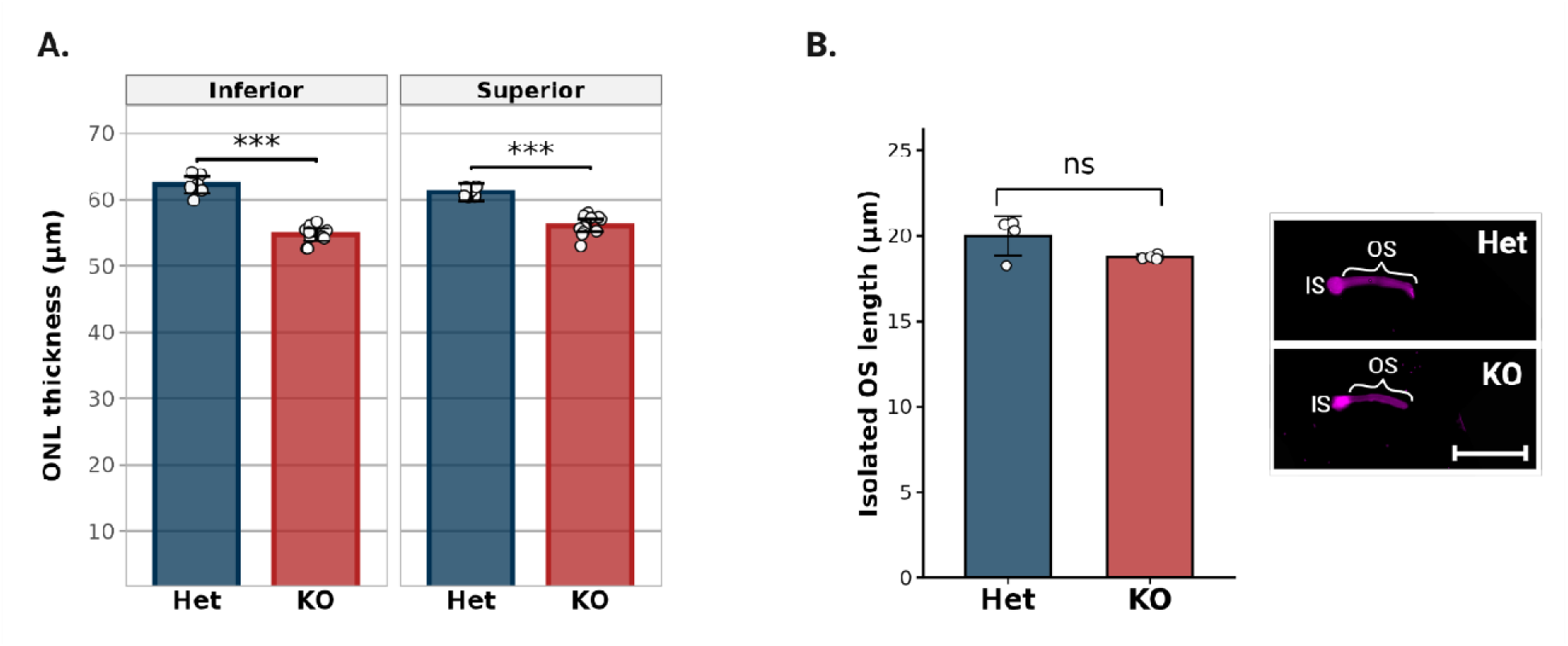
Modest ONL thinning without significant rod outer segment shortening in Kv8.2 KO retina. (**A**) ONL thickness in the inferior and superior retina measured from the same light-adapted OCT images used for the ELM-RPE measurements in Figure 1C. Points represent individual mice; bars show estimated marginal means ± 95% CI (Kv8.2 Het, blue, n = 6; KO, red, n = 11; ***p < 0.001). (**B**) Quantification of mechanically isolated rod OS length. Fifty intact outer segments were measured per mouse and averaged to obtain each animal-level mean. Points represent individual mice; bars show mean ± SD (Kv8.2 Het, blue, n = 4; KO, red, n = 4; p = 0.125, Welch’s t-test). Representative WGA-Alexa 594-labeled outer segments from Kv8.2 Het and KO retina are shown to the right. Scale bar, 20 µm. Abbreviations: ONL, outer nuclear layer; OS, outer segment; IS, inner segment.

To assess the dynamic light-evoked response while minimizing the contribution of any unaccounted for static structural differences, we calculated the light-dark difference in ELM-RPE thickness within each animal (ΔELM-RPE). In the inferior retina, ΔELM-RPE was 2.75 µm lower in Kv8.2 KO mice than in Het controls (95% CI [1.38, 4.11], p < 0.001). A similar reduction was observed in the superior retina (mean difference 2.20 µm, 95% CI [0.84, 3.56], p < 0.01; Fig. 3A; Table S2C). Thus, although ELM-RPE thickness increased with light adaptation in Kv8.2 KO mice, the light-evoked response was attenuated, compatible with changes in SRS fluid handling.

**Figure 3.**
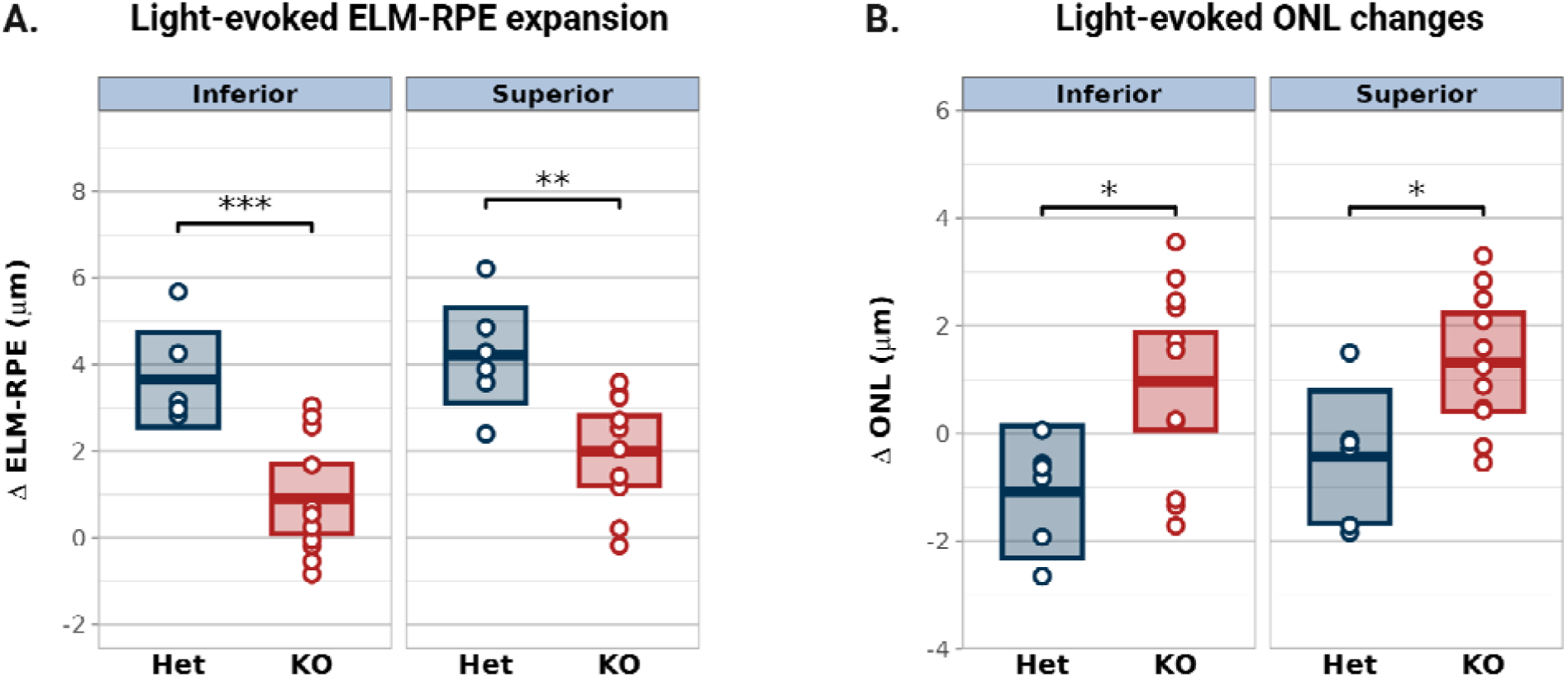
Differential light-evoked changes in ELM-RPE and ONL thickness in Kv8.2 KO retina. (**A**) Light-dark difference in ELM-RPE thickness (ΔELM-RPE) and (**B**) ONL thickness (ΔONL) measured in inferior and superior retina. Measurements are from the same mice and OCT images used in Figure 1C. Points represent individual mice; floating bars show estimated marginal means ± 95% CI (Het, blue, n = 6; KO, red, n = 11; *p < 0.05, **p < 0.01, ***p < 0.001).

We also calculated the light-dark difference in ONL thickness (ΔONL). In the inferior retina, ΔONL was 2.06 µm greater in Kv8.2 KO mice than in Het controls (95% CI [0.53, 3.59], p = 0.0108). A similar genotype difference was observed in the superior retina (mean difference 1.76 µm, 95% CI [0.23, 3.28], p = 0.0264; Fig. 3B; Table S4C). In both regions, the positive mean ΔONL in Kv8.2 KO mice contrasted with the negative mean ΔONL in Het controls, indicating opposing light-dependent changes in ONL and ELM-RPE thickness.

### Retinal molecular changes consistent with an osmosensitive response in Kv8.2 KO retina

We then assessed retinal gene expression by RNA-seq. At four weeks of age, 1,350 genes were differentially expressed in Kv8.2 KO compared with WT retina at adjusted p < 0.01, including 806 increased and 544 decreased genes (Fig. 4A). Pathway analyses showed that genes increased in Kv8.2 KO retina were enriched for developmental, cell-junction and adhesion, extracellular-matrix, and immune-signaling processes, whereas decreased genes were strongly enriched for oxidative phosphorylation, respiratory-chain complexes, and mitochondrial respiration. A curated Müller glia gliosis panel provided evidence of an early reactive response (Supplemental File S2), which is consistent with the increased GFAP protein expression previously observed in older Kv8.2 KO retina.^7^

**Figure 4.**
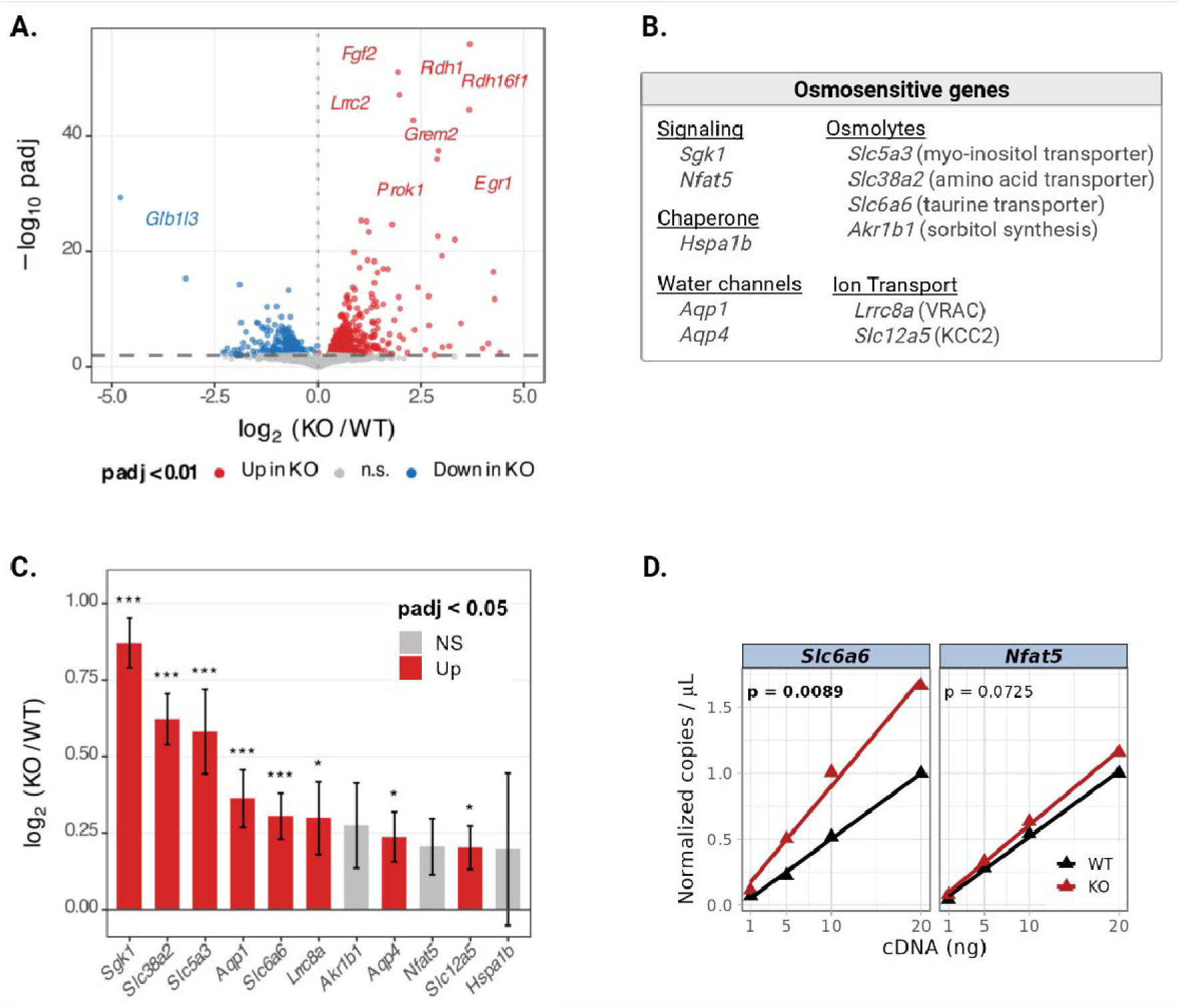
Upregulation of osmosensitive genes in Kv8.2 KO retina. (**A**) Volcano plot of differential gene expression in four-week-old Kv8.2 KO (n = 4) and WT (n = 3) retinal RNA-seq samples. Each point represents one gene plotted by log_₂_ fold change (KO/WT) and −log_₁₀_ adjusted p value. Genes with adjusted p < 0.01 are shown in red (increased in KO) or blue (decreased in KO); genes not meeting this threshold are shown in gray. (**B**) Literature-based panel of osmosensitive genes grouped by functional category. (**C**) Log_2_ fold changes for the genes shown in (B); error bars indicate the uncertainty of each estimate (posterior SD). Red bars indicate genes significantly increased in Kv8.2 KO retina (adjusted p < 0.05), and gray bars indicate nonsignificant genes. Asterisks indicate adjusted p values (*p < 0.05; **p < 0.01; ***p < 0.001). (**D**) RT-ddPCR analysis of *Slc6a6* and *Nfat5* transcript abundance. Points show measurements at individual input amounts of cDNA pooled within genotype, and lines show linear regression fits. Displayed p values test the cDNA input × genotype interaction.

To relate these broad transcriptional changes to the OCT phenotype, we asked whether the RNA-seq dataset contained evidence of an osmosensitive response. Because osmotic stress and cell-volume regulation have been little studied in the retina, we curated a focused, literature-based panel using findings from other tissues (Fig. 4B).^32–42^ The panel included the tonicity-responsive transcription factor NFAT5 and genes reported to respond to NFAT5 or tonicity that function in osmolyte transport or synthesis and molecular chaperoning including *Aqp1* and *Aqp4*, which encode aquaporins abundant in the retina; and *Lrrc8a* and *Slc12a5*, which encode ion-transport proteins involved in regulatory volume decrease (*Lrrc8A* encodes an essential subunit of volume-regulated anion channels (VRAC) and *Slc12a5* encodes a K^+^-Cl^-^ symporter (KCC2)). All 11 genes had positive log₂ fold change estimates in Kv8.2 KO retina, including eight that were significantly increased: five with adjusted p < 0.01 and three additional genes with adjusted p < 0.05 (Fig. 4C). The coordinated increases across multiple functional categories were consistent with an osmosensitive transcriptional response.

We used RT-ddPCR to independently quantify expression of two genes: *Slc6a6*, which encodes the transporter for taurine, one of the most abundant osmolytes in the retina, and *Nfat5*. *Slc6a6* expression was increased in Kv8.2 KO retina, whereas *Nfat5* expression was unchanged, consistent with the RNA-seq results (Fig. 4D).

We next examined aquaporins, which mediate bidirectional water transport across cell membranes. Both *Aqp1*, abundant in photoreceptors, and *Aqp4*, abundant in Müller glia, were upregulated in the RNA-seq dataset. At the protein level, western blotting showed a twofold increase in AQP1 in Kv8.2 KO, but no significant change in AQP4 after normalization to total protein loading (mean difference in AQP1 was 0.71 arbitrary units, 95% CI [0.15, 1.26], p = 0.0284 and mean difference in AQP4 was 0.23, 95% CI [-0.58, 1.01], p = 0.1118; Fig. 5A). A similar increase in labeling intensity for AQP1 in Kv8.2 KO photoreceptor inner segments was also observed by immunolabeling (mean difference 0.77 arbitrary units, 95% CI [0.50, 1.03], p = 0.0065; Fig. 5B). Together, these findings indicate that loss of Kv8.2 is accompanied by increased AQP1 expression.

**Figure 5.**
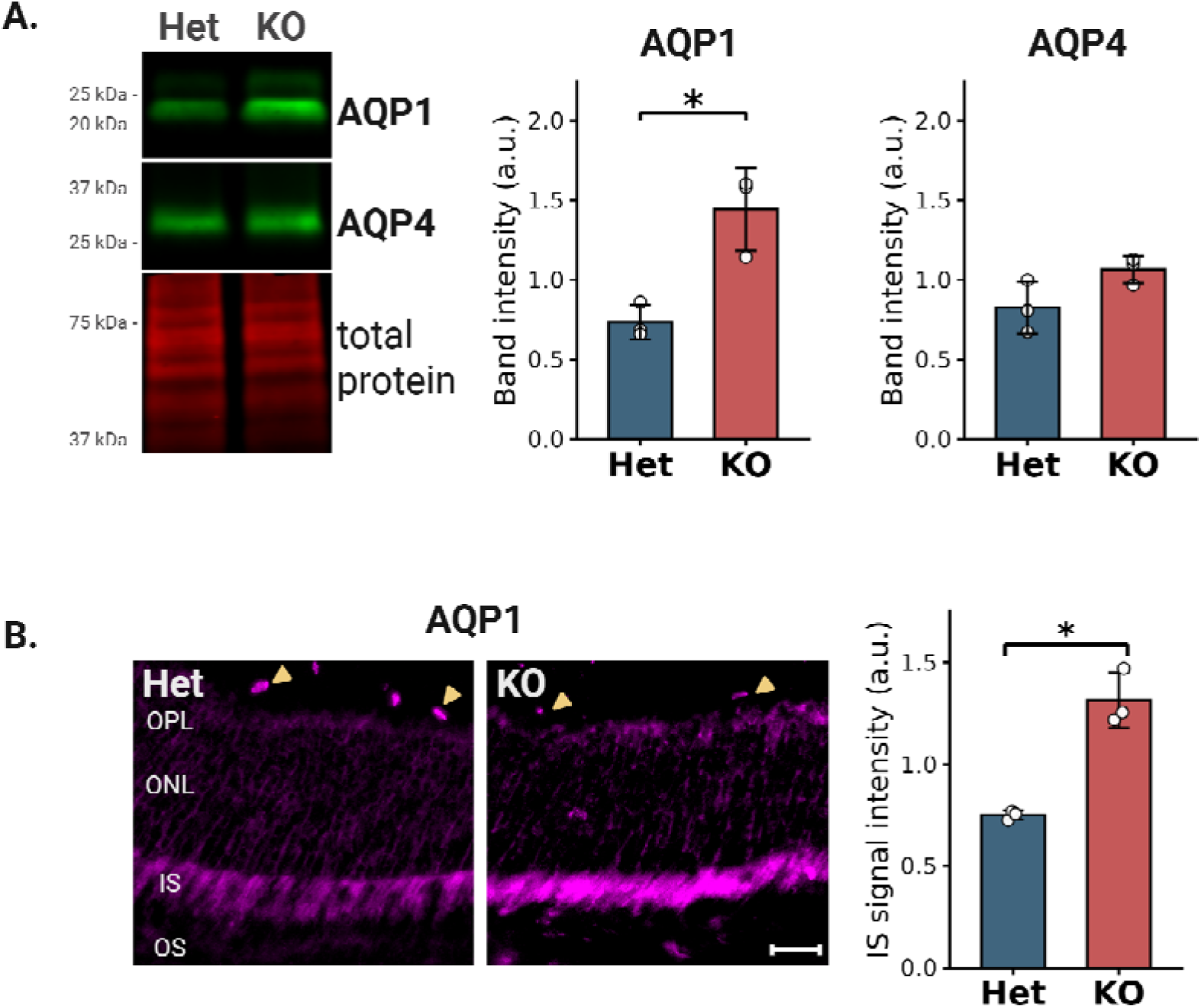
Increased AQP1 protein expression in Kv8.2 KO retina. (**A**) Representative western blots and quantification of AQP1 and AQP4 protein levels in retina from Kv8.2 Het and KO mice. Band intensity was normalized to total protein loading and plotted in arbitrary units (a.u.). (**B**) Immunolabeling of the outer retina with AQP1. Yellow triangles mark presumed red blood cells expressing AQP1. Abbreviations: OS, outer segment, IS, inner segment, ONL, outer nuclear layer, and OPL, outer plexiform layer. Scale bar, 20 µm. Normalized signal intensity per area in the inner segments is plotted in arbitrary units (a.u.). Points represent individual mice; bars show mean ± SD (Het, n = 3; KO, n = 3; *p < 0.05, Welch’s t-test).

## Discussion

We provide noninvasive OCT biomarker and molecular evidence consistent with altered fluid homeostasis in the outer retina following loss of Kv8.2. Specifically, Kv8.2 KO retina exhibited i) reduced light-dependent ELM-RPE expansion, compatible with changes in SRS fluid handling; ii) a light-dependent expansion of the outer nuclear layer; iii) an osmosensitive transcriptional response; and iv) increased expression of aquaporin-1 in photoreceptors. Together, these findings support a model in which potassium efflux through Kv2.1/Kv8.2 channels contributes to fluid distribution between photoreceptors and the SRS.

In the retina, coupling of potassium flux to water movement is already well established in Müller glia, where inwardly rectifying Kir4.1 channels and AQP4 coordinate ion and fluid transport.^43^ By analogy, a similar relationship may exist in photoreceptors, where Kv2.1/Kv8.2 channels and AQP1 are positioned at the inner segment plasma membrane adjacent to the SRS. In this context, potassium efflux through Kv2.1/Kv8.2 channels–best known for shaping the photoreceptor light response–could contribute to local osmotic gradients that facilitate water movement from photoreceptors into the SRS for subsequent clearance across the RPE (Fig. 6).

**Figure 6.**
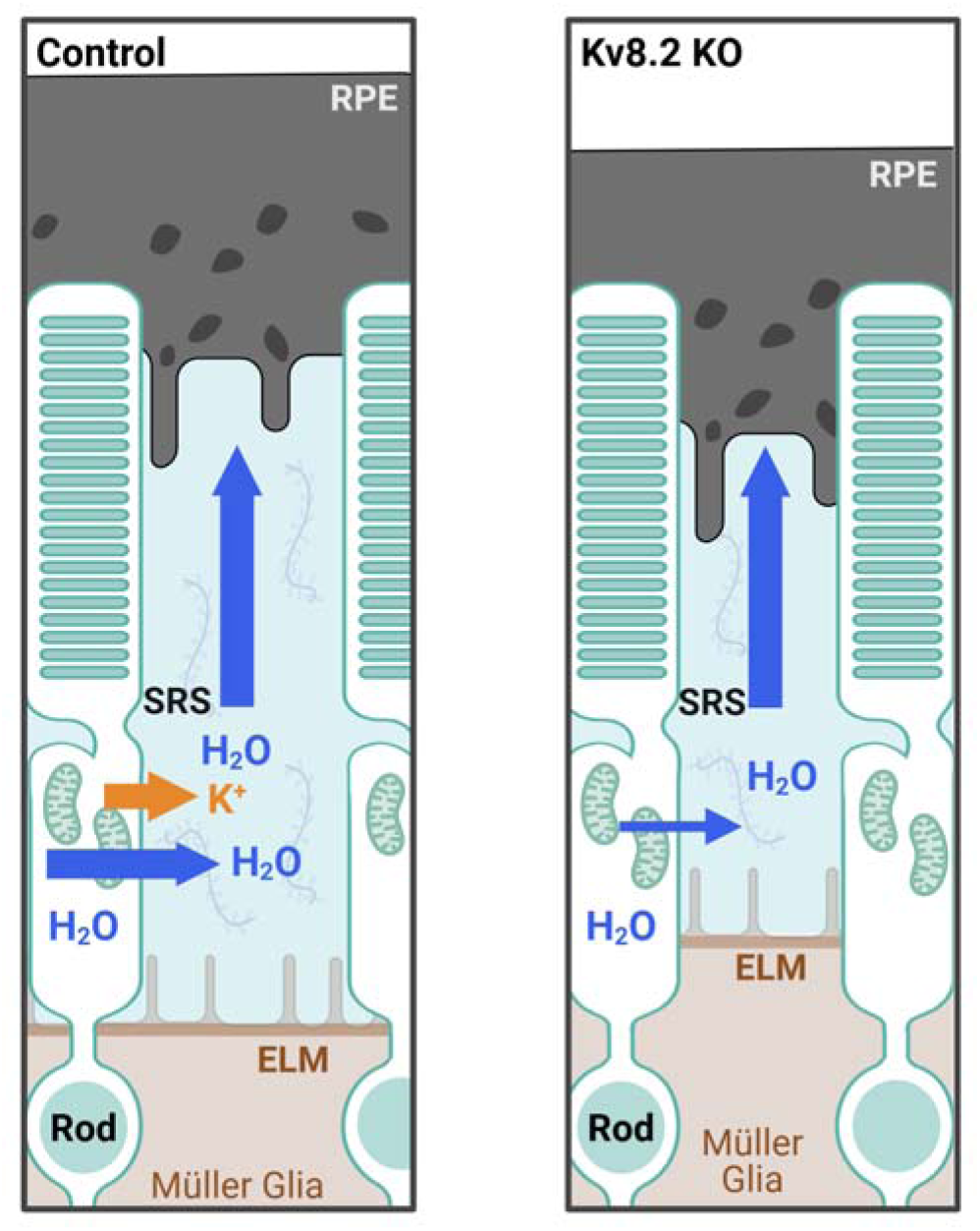
Proposed model linking Kv2.1/Kv8.2 mediated potassium flux to osmotic regulation in the outer retina. Schematic comparing control (left) and Kv8.2 KO (right) retina. The SRS extends from the ELM to the RPE and surrounds the photoreceptor inner and outer segments. Water moves from photoreceptors through the SRS toward the RPE (blue arrows), while Kv2.1/Kv8.2 channels mediate KD efflux from photoreceptors (orange arrow). We hypothesize that this KD efflux contributes to an osmotic gradient that promotes water movement into the SRS. Loss of functional Kv2.1/Kv8.2 channels is therefore predicted to reduce KD and water efflux from photoreceptors. Assuming water uptake by the RPE remains unchanged, this imbalance is predicted to reduce SRS volume, consistent with the attenuated light-evoked ELM-RPE expansion measured by OCT in Kv8.2 KO retina.

The ELM-RPE metric is an anatomical measurement, but it is also a biomarker of dynamic outer retina physiological processes. We considered that since Kv8.2 KO undergoes a slow retinal degeneration potential OS shortening could influence the changes in ELM-RPE that we found. Two observations inform our interpretation of the reduced light-evoked ELM-RPE expansion in Kv8.2 KO retina. First, despite modest ONL thinning, we detected no significant shortening of rod outer segments, the structural component within the ELM-RPE interval most likely to be affected by retinal degeneration. Second, the paired light-dark difference (ΔELM-RPE) minimizes contributions from fixed genotype-specific anatomical differences and identifies an altered light-evoked response compatible with changes in SRS fluid handling. Because loss of Kv2.1/Kv8.2 channels depolarizes photoreceptors and alters their light response, the attenuated ELM-RPE response could also arise indirectly through changes in photoreceptor metabolic demand, SRS acidification, or pH-dependent fluid clearance by the RPE, in addition to a direct effect of reduced potassium efflux on osmotic water movement.^18,27^ The present experiments cannot distinguish between these proposed mechanisms or exclude contributions from other light-response changes.

The observation of the light-dark difference in ONL thickness that we observed in Kv8.2 KO mice is novel to the best of our knowledge. Prior work using acetazolamide has shown that acute perturbation of retinal fluid handling can reduce ONL thickness in light-adapted retina, supporting a link between ONL thickness and hydration.^44^ In the present study, ΔONL increased in Kv8.2 KO while ΔELM-RPE decreased, raising the possibility that relative expansion of the ONL reflects impaired redistribution of fluid from photoreceptors into the SRS, resulting in water retention within photoreceptors. Alternatively, the expansion could reflect structural remodeling or fluid accumulation in Müller glia. Additional studies will be required to define the physiological basis of this phenomenon.

Four limitations should be considered when interpreting the molecular findings. First, our use of bulk neural retina does not allow us to identify the cellular source of the response.

Photoreceptors are one likely contributor because they are the most abundant retinal cell type, express Kv2.1/Kv8.2 channels, and have increased aquaporin-1 expression in the Kv8.2 KO. However, the early reactive Müller glia response identified by RNA-seq and the established role of Müller glia in retinal fluid homeostasis suggest that these cells may also contribute. Second, our RNA-seq analysis did not examine the RPE, which was removed during neural-retina dissection. The RPE is a major determinant of SRS fluid volume through coordinated ion and water transport^15^. Altered RPE responses may therefore contribute to the Kv8.2 KO phenotype but were not evaluated in this study. Third, regulation of osmolyte transporters and aquaporins occurs through mechanisms in addition to gene expression, including trafficking and post-translational modification.^45–49^ Finally, the structural and molecular analyses were performed in separate cohorts of young mice at similar but different ages (RNA-seq, 4 weeks; OCT, ∼5 weeks (P33-P41); protein analysis, 8 weeks; and RT-ddPCR, 14 weeks). Although selected molecular changes remained detectable at later ages, these experiments do not define their temporal relationship to the OCT phenotype. Consequently, the changes reported here should not be interpreted as direct evidence for activation of a specific tonicity signaling pathway, but rather as evidence that loss of Kv8.2 is accompanied by a coordinated molecular response involving osmolyte and water homeostasis.

In summary, the convergence of structural and molecular changes support a model in which potassium efflux through Kv2.1/Kv8.2 channels contributes to regulation of fluid balance in the outer retina. Altered outer retinal fluid regulation may represent an underappreciated component of KCNV2 retinopathy and could have broader relevance for retinal diseases in which ion fluxes are altered.

## Acknowledgments

We thank the NIH/NEI Center Support Grant to the University of Iowa (P30EY025580) for provision and maintenance of the Bioptigen spectral-domain OCT imaging system.

## Commercial Relationships Disclosures

None

