## Supplemental for "Biomarkers indicate altered outer retinal fluid homeostasis in the absence of Kv2.1/Kv8.2 channels"

### Index of Supplemental material

Figure S1: Validation of Kv8.2 Het as control, ERG. Related to Methods.

Figure S2: Validation of Kv8.2 Het as control, OCT. Related to Methods.

Figure S3: Uncropped western blots. Related to Figure 5

File S1: Statistical outputs for OCT data (Tables S1A-S4C). Related to Figures 1-3.

File S2: RNA-seq bioinformatic analysis and supporting results. Related to Figure 4.

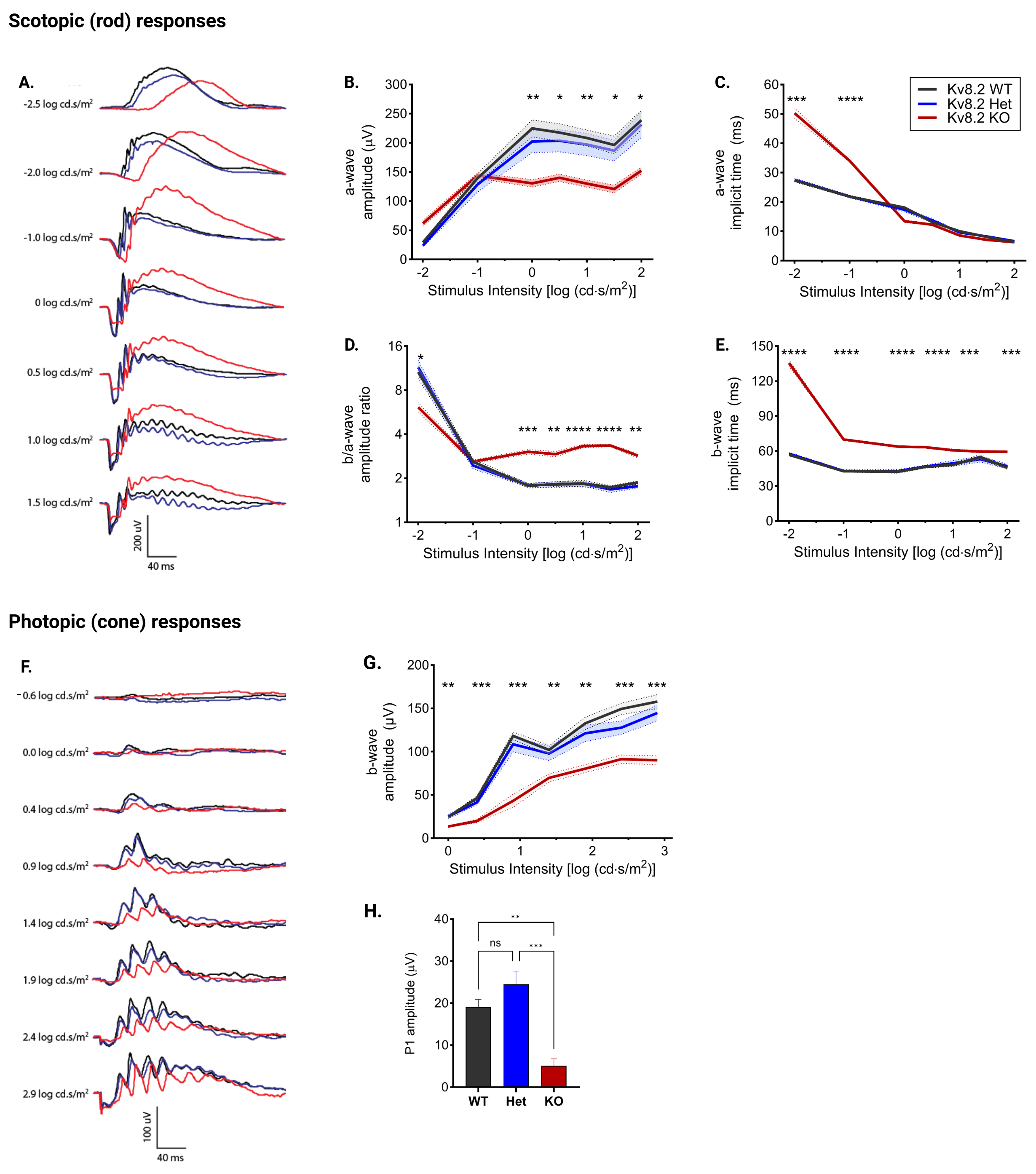

### Supplemental Figure S1. Kv8.2 Het ERG responses were indistinguishable from WT.

To reduce the number of animals required for this study we first assessed whether Kv8.2 heterozygous mice could serve as littermate controls using ERG as previously described.^7^ Five Kv8.2 WT (black), seven Het (blue), and four KO (red) animals were used for the intensity series and five of each genotype for the 30 Hz flicker were used. The cohorts included both sexes and animals were two-months old. ERG data was analyzed using 2-way repeated measures ANOVA followed by Tukey's multiple comparisons test or 1-way ANOVA for panel H (GraphPad Prism, Ver. 10, statistical data tables available on request).

Panels A and F show representative scotopic and photopic ERG traces, respectively, from WT (black), Het (blue), and KO (red) mice. The characteristic Kv8.2 KO rod driven responses were observed: an a-wave with normal amplitude but increased time to peak at low light intensity that switched to reduced amplitude with normal timing at high light intensity (**A-C**); subnormal b/a-wave amplitude at the dimmest light intensities that switched to supernormal at higher light intensities (**D**); and delayed time to peak for the b-wave at all light intensities (**E**). Under light-adapted conditions, cone-driven responses were evoked by an intensity series of bright flashes (**F-G**) or a 30 Hz flicker (**H**). As expected, cone responses from Kv8.2 KO mice had a reduced b-wave amplitude to single flashes and a diminished P1 amplitude to high-frequency flicker stimuli. Quantitative data in panels B-E, G, and H are plotted as mean ± SEM. In panels B-E, G, the line representing mean often obscures the narrow SEM error bands, asterisks above given light intensities indicate statistically significant differences between WT and KO (p < 0.05 (*); p < 0.01 (**); p<0.001 (***); p < 0.0001 (****)). There was no statistically significant difference between Kv8.2 WT and Het rod or cone responses.

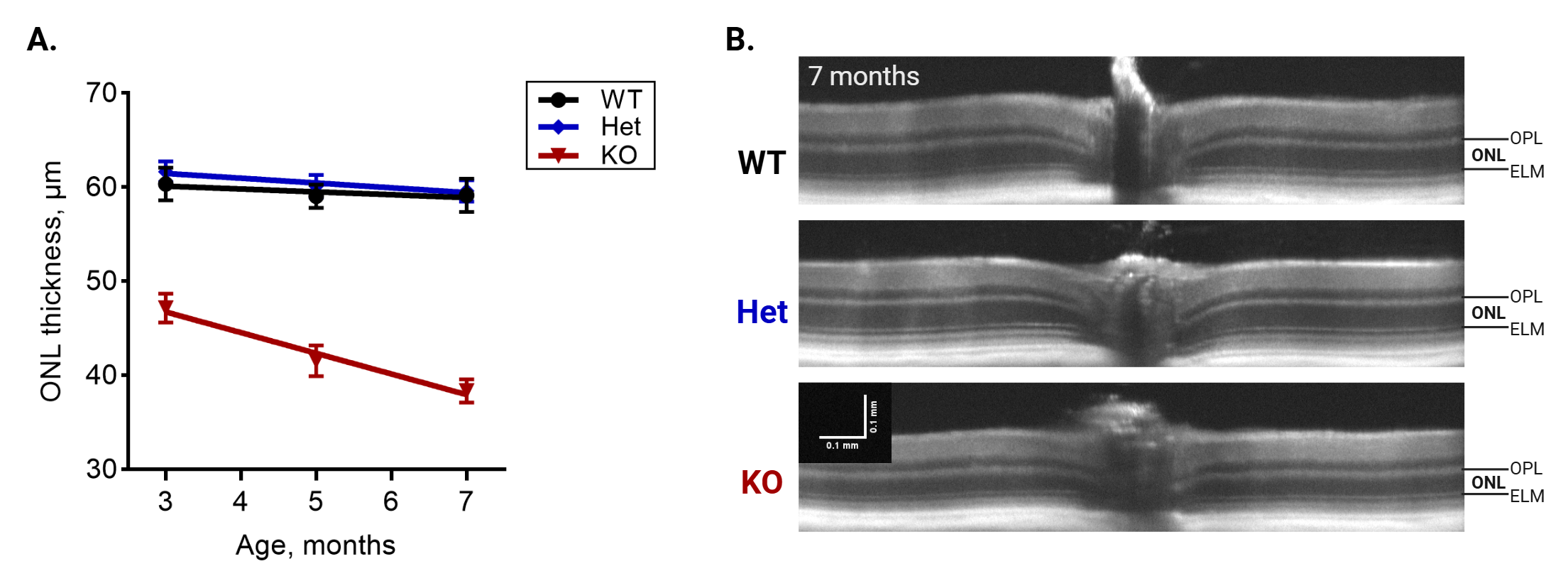

### Supplemental Figure S2. ONL thinning over time was not present in Kv8.2 Het.

ONL thickness was measured by OCT as described in the main text for a cohort of 4 Kv8.2 WT (black), 10 Kv8.2 Het (blue), and 5 Kv8.2 KO (red) light-adapted mice at 3, 5, and 7 months of age to check for retinal degeneration in Kv8.2 Hets. Data are plotted as mean ± SD, with fitted linear regression lines (**A**). Representative images for each genotype are shown at 7 months are shown in (**B**); horizontal and vertical scale bars, 0.1 mm. Consistent with our prior report using manual measurement of ONL thickness, the ONL in Kv8.2 KO mice declined modestly (20–36%) from 3–7 months of age, whereas ONL thickness remained stable in both WT and Het retinas. Together with Supplemental Figure 1, this data confirms that either Kv8.2 Het or Kv8.2 WT may be used as controls for Kv8.2 KO littermates.

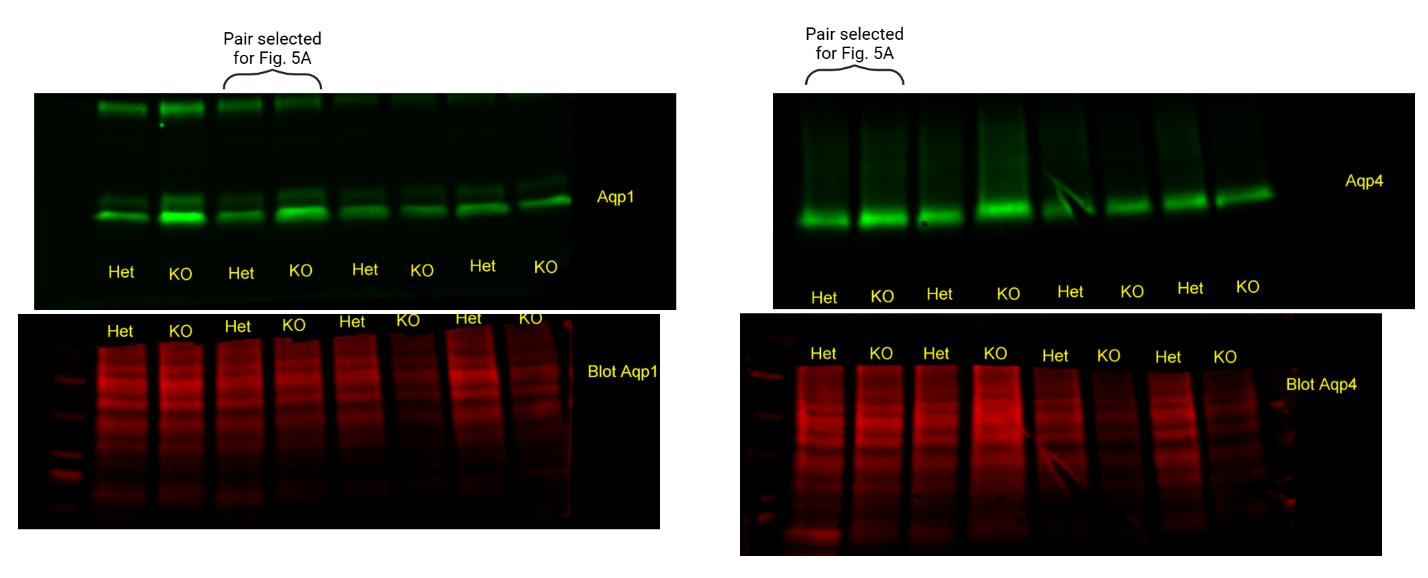

### Supplemental Figure S3. Uncropped, unedited western blots corresponding to the representative images in Figure 5A. Lanes 1-4 and 5-8 contain technical duplicates of samples from two of the biological replicates per genotype included in the quantification.
