## Supplemental File S2 for "Biomarkers indicate altered outer retinal fluid homeostasis in the absence of Kv2.1/Kv8.2 channels"

### Mouse retinas: KCNV2-KO vs WT RNA-seq Analysis

Revision 9 — KO2 excluded on knockout validation, SVA-adjusted

AUTHOR
Miles Pufall

PUBLISHED
July 31, 2026

#### Revision history

| File | Change |
|---|---|
| `sheila_rnaseq_KOvWT_3.qmd` | Original. KO2 and KO3 dropped on QC grounds (leaving 3 WT vs 3 KO of an original 3 vs 5); unshrunken log2FC; volcano at padj < 0.05 with a 20% FC cutoff. |
| `sheila_rnaseq_KOvWT_4.qmd` | All eight samples retained. Every analysis run twice, ± SVA, to test whether the adjustment was absorbing genotype signal. |
| `sheila_rnaseq_KOvWT_5.qmd` | Streamlined to the SVA-adjusted model throughout. Added the two-tier Müller glia gliosis panel and an over-representation suite (GO, KEGG, Reactome, WikiPathways) alongside GSEA; ± SVA comparison condensed to an appendix. |
| `sheila_rnaseq_KOvWT_6.qmd` | Gene-panel bar charts at padj < 0.05; osmosensitive panel restored; shrunken estimates reported plainly as `log2FC`. |
| `sheila_rnaseq_KOvWT_8.qmd` | *Kcnv2* knockout validation added as an explicit pre-modelling QC step; **KO2 excluded** because it shows no knockdown. KO3 retained. Dataset rebuilt, renormalised and SVA re-estimated on the 7 retained samples. Axis clipping to ±5 applies to the volcano panels only. |

`sheila_rnaseq_KOvWT_9.qmd` | **This file.** Input paths validated up front via a `project_dir` setting, so a missing `quants/` fails immediately with a readable message instead of silently producing an empty sample table. |

#### Design and rationale

Eight libraries were sequenced: **WT1–3 vs KO2–6** (3 WT, 5 KO). Revision 3 dropped KO2 and KO3 because they separated in the PCA; that decision is not carried forward, since removing samples on the basis of a PCA computed from the same expression data is circular.

**KO2 is excluded here on a different and non-circular basis:** it shows no reduction in *Kcnv2* itself, so there is no evidence the knockout is present in that animal. The analysis therefore runs on **3 WT vs 4 KO**. KO3 is retained — it separated in the original PCA but its *Kcnv2* level sits with the true knockouts.

The structure those two samples create is instead **modeled**: surrogate variable analysis estimates it and it enters the design as a covariate. The appendix documents why this is defensible here — SV1 carries only $R^2 \approx 0.12$ with genotype, and the fold changes of the strongly-called genes are essentially unchanged with or without the adjustment, so the SV is absorbing technical structure rather than competing with the contrast.

Conventions used throughout:

- **Design:** `~ SV1 (+ SV2) + background`, contrast KO vs WT.
- **Effect sizes:** all reported `log2FC` values are apeglm-shrunken, which is simply the fold change this design supports; the unshrunken maximum- likelihood values are kept as `log2FC_MLE` for reference.
- **Significance:** `padj < 0.01`, no fold-change cutoff.
- **Volcano:** red up in KO, blue down, x-axis clipped to ±5 with off-scale genes counted in the subtitle rather than dropped. This clip applies to the volcano panels alone — every other figure is scaled to its own data.
- **Figure export:** every panel is written to `figures_pub/` as a 4 × 4 cm PDF, drawn at that size rather than scaled down. Legible versions appear inline.

### Trimming and Mapping

#### Adapter trimming with TrimGalore! (v0.6+)

```
for i in *_R1_001.fastq.gz;
do
  SAMPLE=$(echo ${i} | sed "s/_R1_001\.fastq.gz//")
  trim_galore --paired -j 4 ${SAMPLE}_R1_001.fastq.gz ${SAMPLE}_R2_001.fastq.gz
done
```

#### Alignment and quantification with Salmon (v1.10.2)

```
for i in *_R1_val_1.fq.gz;
do
  SAMPLE=$(echo ${i} | sed "s/_R1_val_1\.fq.gz//")
  salmon quant -i
/Shared/pufall/mpufall/RNAseq/DataSets2015/3722_Dex_v_Pred/Trimmed/gen_v45_tx_index/ \
    -p 24 -l A -1 ${SAMPLE}_R1_val_1.fq.gz -2 ${SAMPLE}_R2_val_2.fq.gz \
    --validateMappings --seqBias --gcBias --posBias --numBootstraps 50 \
    -o quants/${SAMPLE}
done
```

### Setup

▶ Code

#### Constants and the publication figure system

▶ Code

### Mapping statistics

▶ Code

```
Found 8 samples in
/Users/mpufall/Library/CloudStorage/Dropbox/miles/projects/sheila/rnaseq/quants
```

▶ Code

Samples included in this analysis

| names | background | rep |
|---|---|---|
| WT1 | WT | 1 |
| WT2 | WT | 2 |
| WT3 | WT | 3 |
| KO2 | KO | 2 |
| KO3 | KO | 3 |
| KO4 | KO | 4 |
| KO5 | KO | 5 |
| KO6 | KO | 6 |

▶ Code

```
Parsed fields: salmon_version, samp_type, opt_type, num_libraries, library_types,
frag_dist_length, frag_length_mean, frag_length_sd, seq_bias_correct, gc_bias_correct,
num_bias_bins, mapping_type, keep_duplicates, num_valid_targets, num_decoy_targets,
num_eq_classes, serialized_eq_classes, index_seq_hash, index_name_hash,
index_seq_hash512, index_name_hash512, index_decoy_seq_hash, index_decoy_name_hash,
num_bootstraps, num_processed, num_mapped, num_decoy_fragments, num_dovetail_fragments,
num_fragments_filtered_vm, num_alignments_below_threshold_for_mapped_fragments_vm,
percent_mapped, call, start_time, end_time
```

▶ Code

Mean depth 25,454,187 read pairs; mean mapping rate 83%.

▶ Code

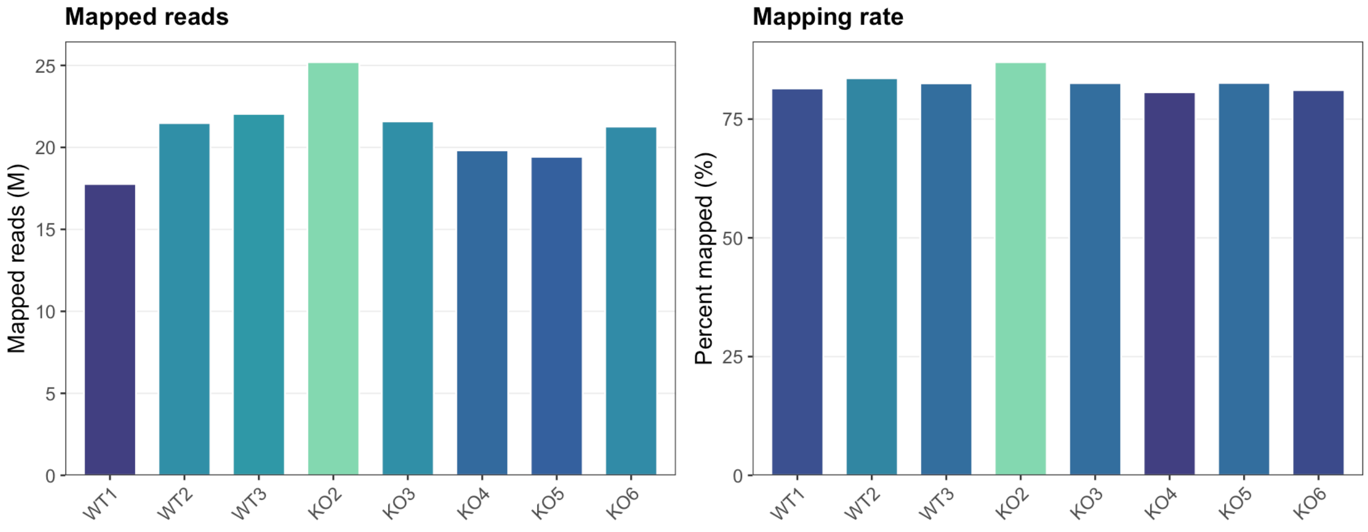


### Import into DESeq2

▶ Code

```
Import genomic features from the file as a GRanges object ... OK
Prepare the 'metadata' data frame ... OK
Make the TxDb object ... OK
```

▶ Code

```
'select()' returned 1:1 mapping between keys and columns
```

▶ Code

```
reading in files with read_tsv
1 2 3 4 5 6 7 8
transcripts missing from tx2gene: 64
summarizing abundance
summarizing counts
summarizing length
```

▶ Code

▶ Code

```
Genes retained after filtering: 20341
```

▶ Code

```
Samples: 8
```

### Sample QC

▶ Code

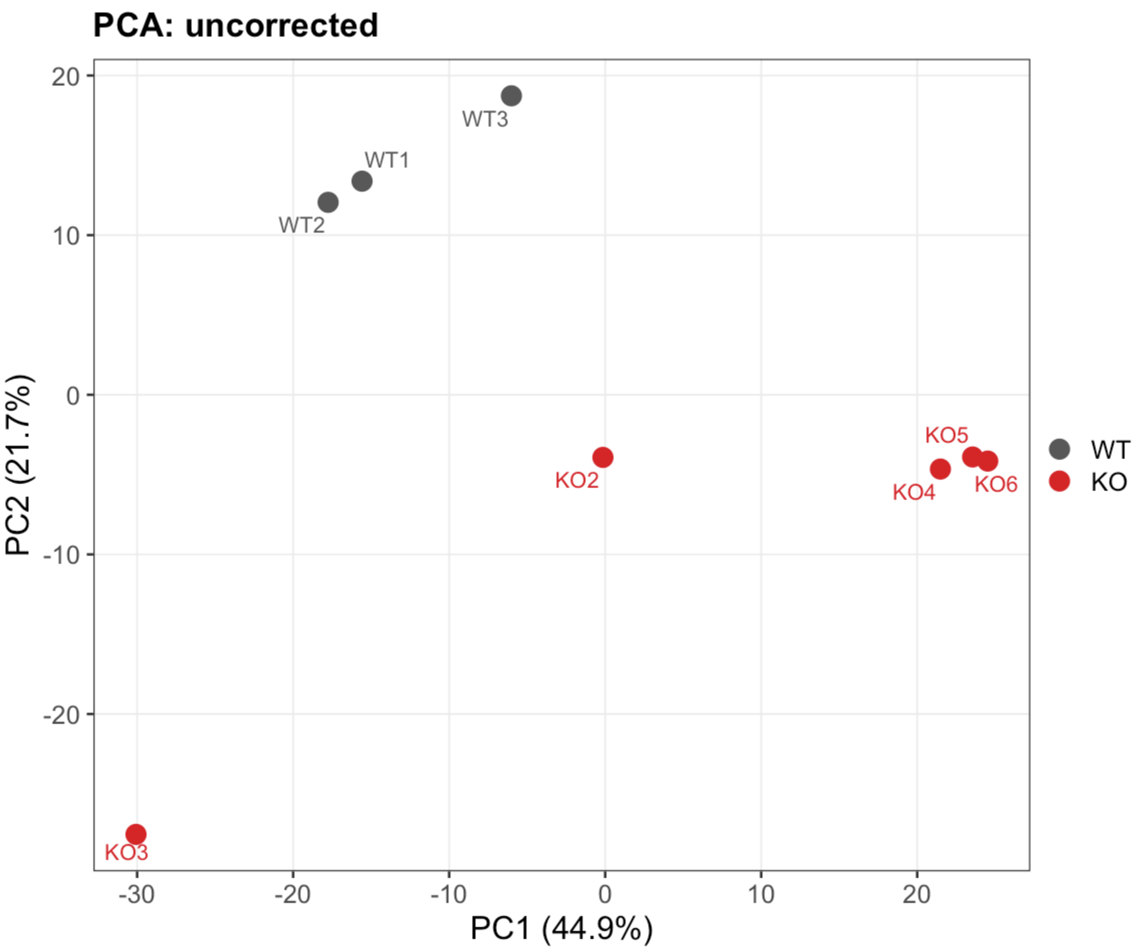
PCA: uncorrected
WT3
WT1
WT2
KO5
KO2
KO4
KO6
KO3
WT
KO
PC2 (21.7%)
PC1 (44.9%)
20
10
0
-10
-20
-30
-20
-10
0
10
20

▶ Code

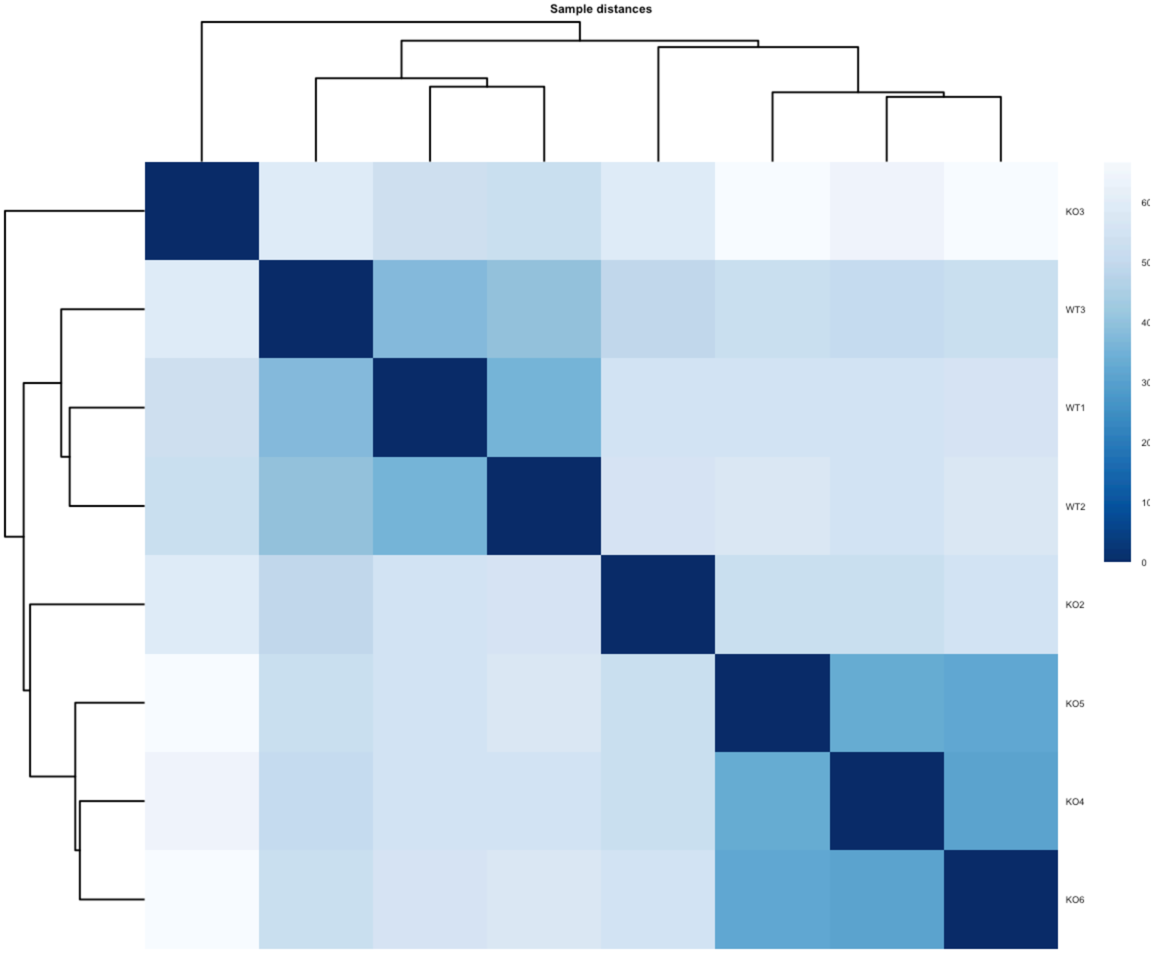


▶ Code

▶ Code

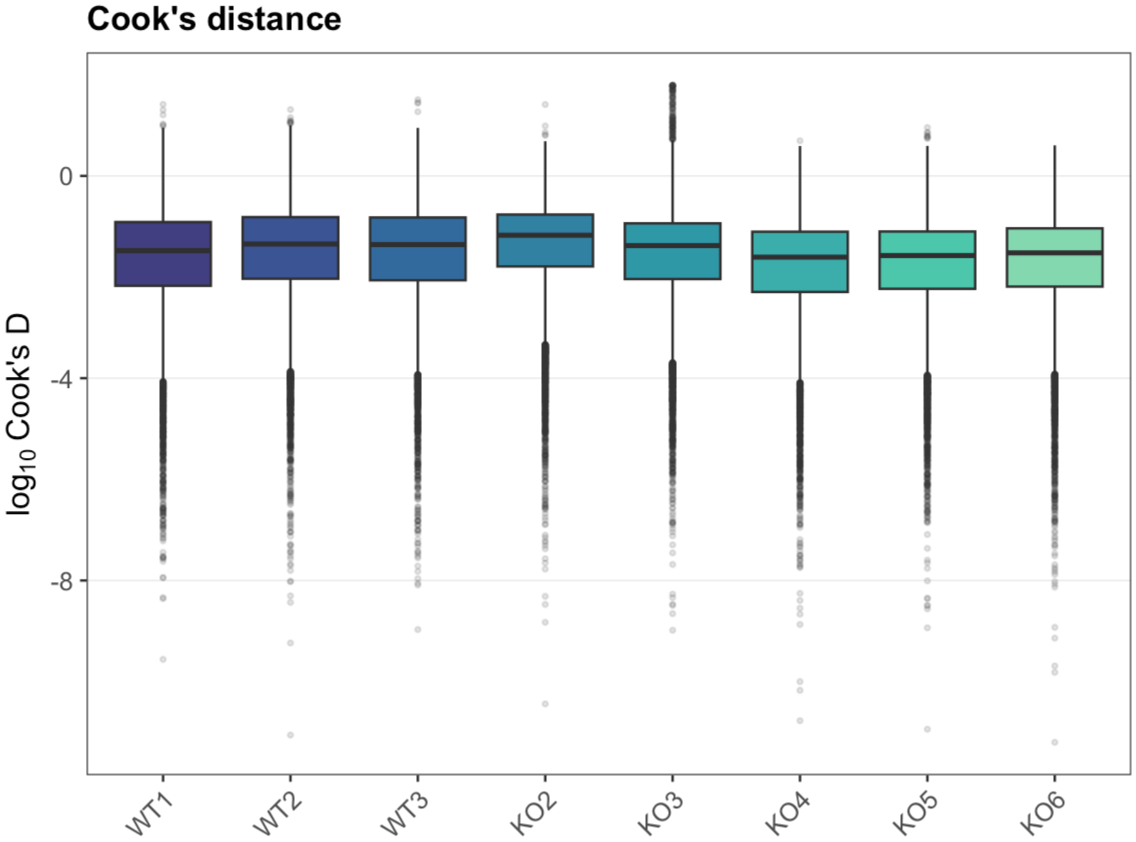
Cook's distance
log10 Cook's D
0
-4
-8
WT1
WT2
WT3
KO2
KO3
KO4
KO5
KO6

▶ Code

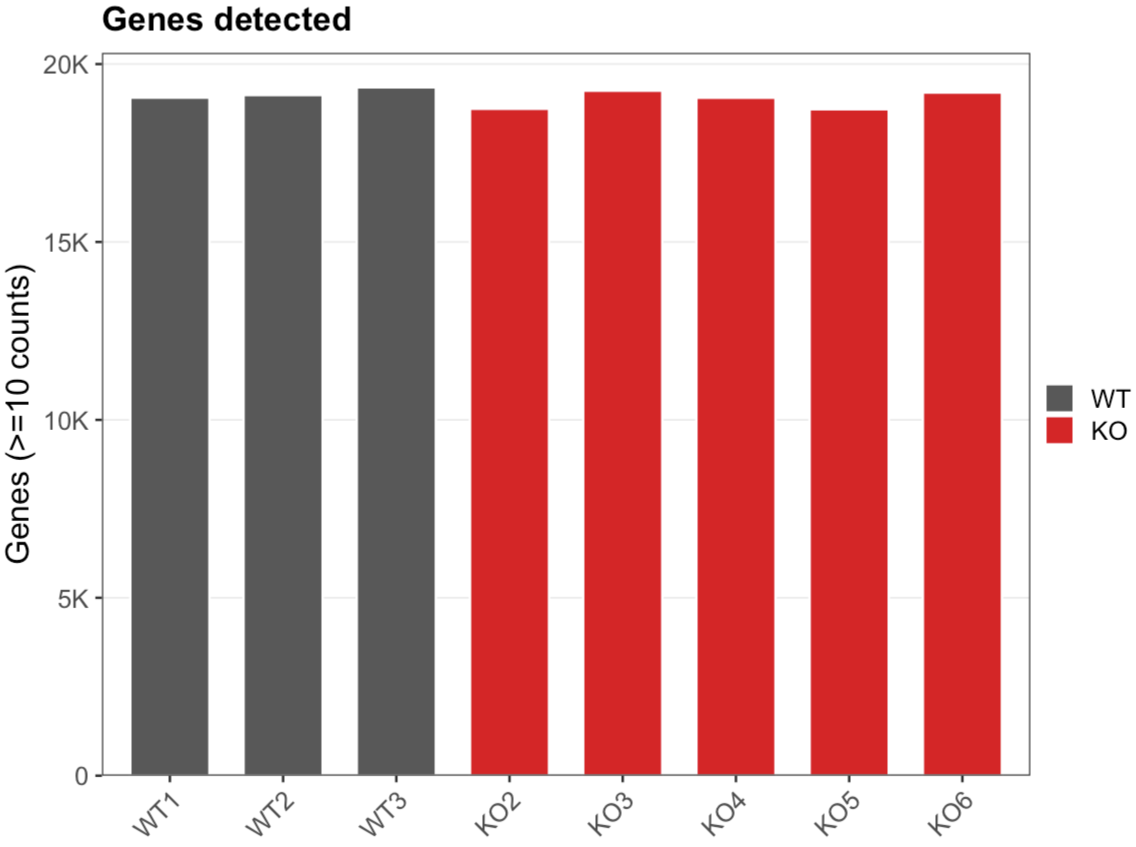
Genes detected
Genes (>=10 counts)
20K
15K
10K
5K
0
WT1
WT2
WT3
KO2
KO3
KO4
KO5
KO6
WT
KO

### Knockout validation and sample exclusion

Before any modelling, confirm the knockout is actually present in each animal labelled KO. This is the one sample-level exclusion criterion in the analysis, and it is deliberately **external to the expression structure**: it asks whether the experimental manipulation worked in a given animal, not whether that animal resembles its group-mates transcriptome-wide. That distinction is what separates it from the PCA-based exclusion in revision 3, which used the data to decide which data to keep.

▶ Code

Kcnv2 normalized counts, all samples

| sample | background | count | pct_of_WT_min | knockdown |
|---|---|---|---|---|
| WT1 | WT | 10574 | 100 | - |
| WT2 | WT | 11232 | 106 | - |
| WT3 | WT | 10976 | 104 | - |
| KO2 | KO | 12458 | 118 | NO |
| KO3 | KO | 9273 | 88 | yes |
| KO4 | KO | 8383 | 79 | yes |
| KO5 | KO | 8729 | 83 | yes |
| KO6 | KO | 8789 | 83 | yes |

▶ Code

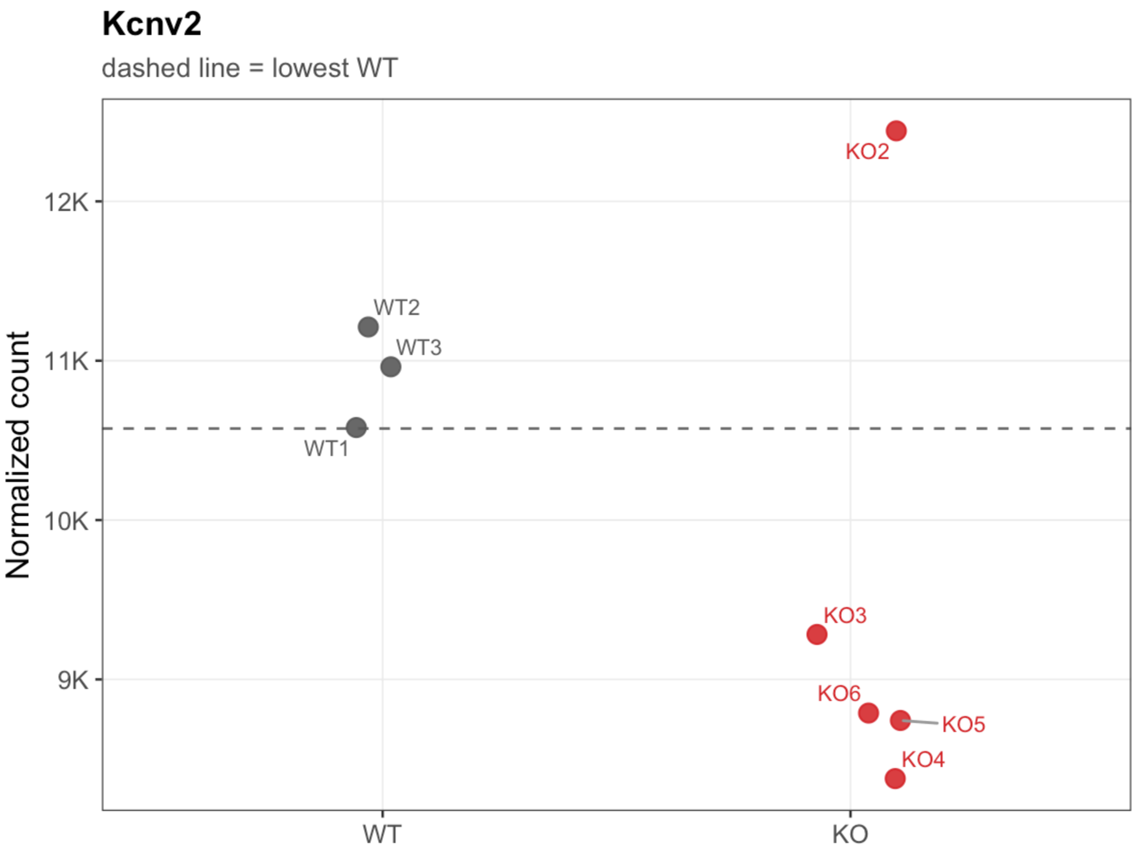


▶ Code

```
KO samples with NO evidence of Kcnv2 knockdown: KO2
```

▶ Code

```
EXCLUDED FROM ALL DOWNSTREAM ANALYSIS: KO2
Reason: no reduction in Kcnv2 relative to the WT range, i.e. the knockout is not
detectable in this animal.
```

▶ Code

```
Retained: 7 samples (3 WT, 4 KO)
```

**Sample exclusion**

**KO2 is excluded from all downstream analysis.** Its *Kcnv2* level is not merely within the WT range but above every WT sample, which is not what an animal carrying the knockout allele should look like. The most likely explanations are a mis-genotyped animal or a sample swap at collection or library preparation.

The exclusion is applied **before** normalisation, filtering, dispersion estimation and surrogate variable analysis, because all four use every sample in the object. Dropping KO2 only at the results stage would leave its influence baked into the size factors and the surrogate variables.

KO3 is **retained**. It separated alongside KO2 in the original PCA, but its *Kcnv2* level sits with the other true knockouts, so the knockout-verification criterion keeps it. Excluding it would require falling back on the PCA argument this re-analysis rejects.

Worth confirming independently by genotyping PCR if DNA remains — a transcript-level result is suggestive of a swap, not proof of one.

##### Rebuilding the dataset on the retained samples

▶ Code

```
Genes retained after filtering: 20133
```

▶ Code

```
Samples in the model: 7
```

#### Surrogate variables

▶ Code

```
SVs suggested by num.sv (be): 1
```

▶ Code

```
SVs used: 1
```

▶ Code

```
Number of significant surrogate variables is:  1
Iteration (out of 5 ):1  2  3  4  5
```

An SV orthogonal to genotype can only remove noise; the more of the genotype contrast it contains, the more real effect it removes alongside. `R²` is from regressing each SV on genotype — near 0 is safe.

▶ Code

Association of each surrogate variable with genotype

| SV | R2 | p_value | cor_geno |
|---|---|---|---|
| SV1 | 0.09 | 0.514 | -0.299 |

▶ Code

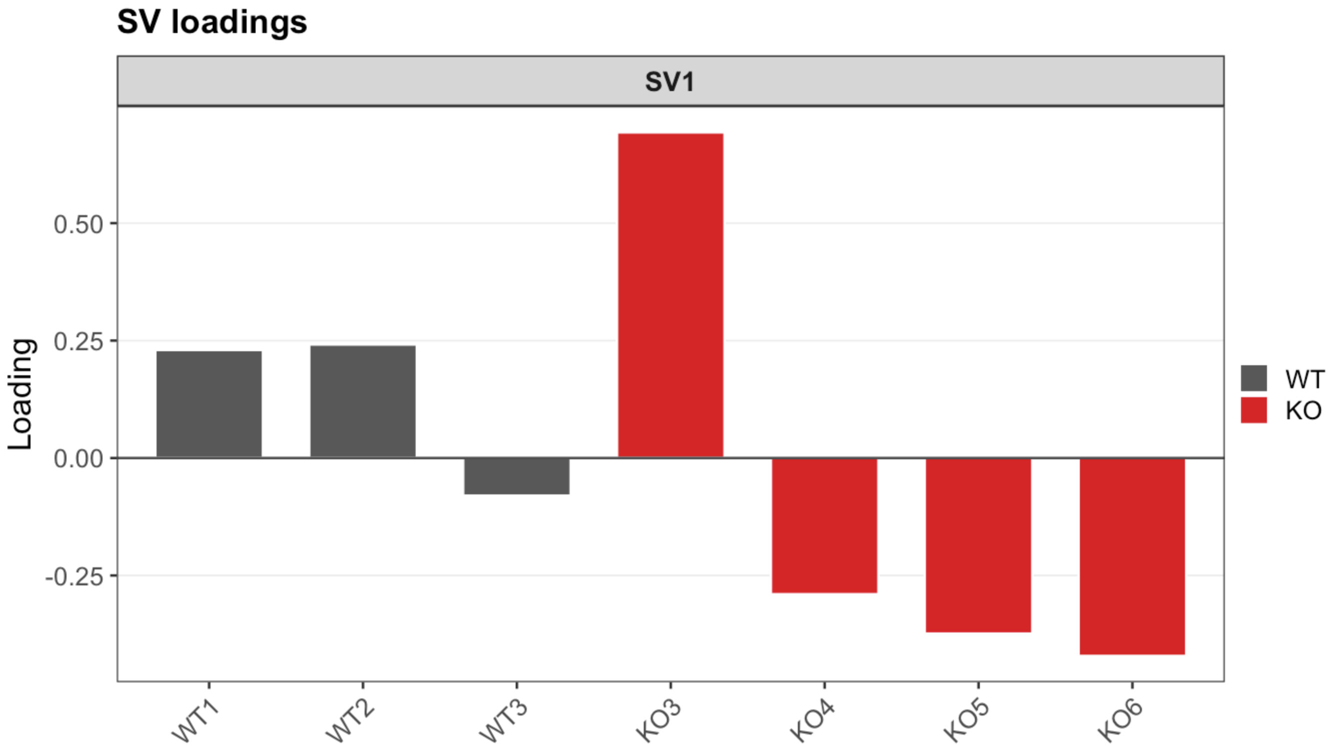


▶ Code

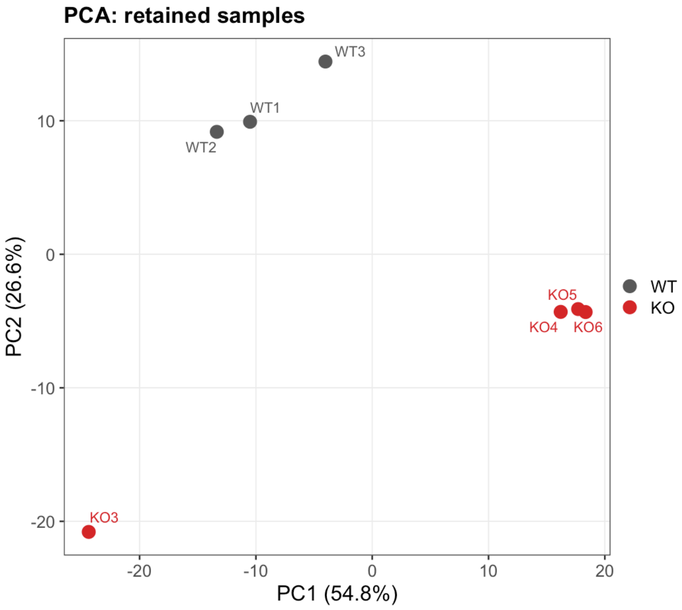


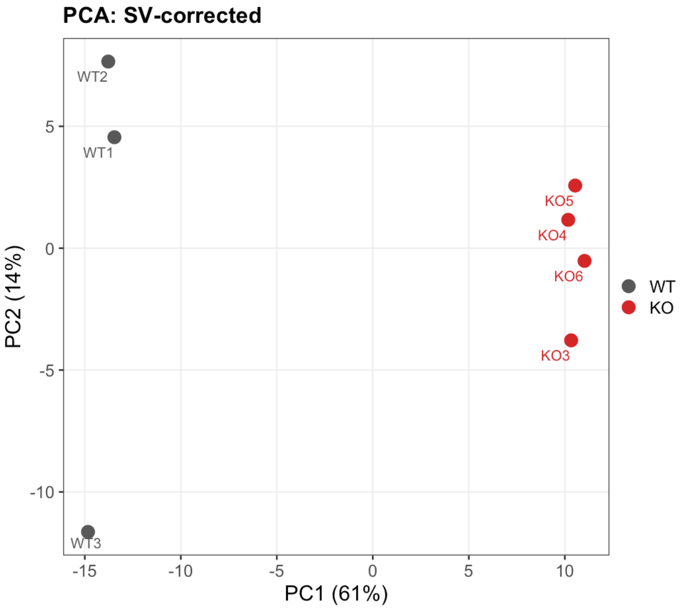


### Differential expression: KO vs WT

▶ Code

```
Design: ~SV1 + background
```

▶ Code

```
out of 20133 with nonzero total read count
adjusted p-value < 0.01
LFC > 0 (up)       : 806, 4%
LFC < 0 (down)     : 544, 2.7%
outliers [1]       : 0, 0%
low counts [2]     : 3123, 16%
(mean count < 20)
[1] see 'cooksCutoff' argument of ?results
[2] see 'independentFiltering' argument of ?results
```

▶ Code

#### Annotation and export

▶ Code

```
Genes with a usable symbol: 20080 of 20133
```

▶ Code

```
Genes with a usable Entrez ID: 15950 of 20133
```

▶ Code

```
Significant at padj < 0.01: 1350 (806 up, 544 down)
```

▶ Code

Top 25 genes by adjusted p-value

| symbol | log2FC | lfcSE | padj | baseMean |
|---|---|---|---|---|
| Vim | 1.499 | 0.078 | 0 | 2804.635 |
| Fgf2 | 1.967 | 0.108 | 0 | 2405.789 |
| Lrrc2 | 2.003 | 0.119 | 0 | 2055.323 |
| mt-Atp8 | -1.759 | 0.111 | 0 | 28256.884 |
| Grem2 | 2.240 | 0.142 | 0 | 497.227 |
| Krt18 | 1.647 | 0.108 | 0 | 1506.988 |
| Rdh1 | 3.643 | 0.239 | 0 | 211.381 |
| Gckr | 1.492 | 0.099 | 0 | 1628.327 |
| mt-Nd6 | -1.362 | 0.090 | 0 | 27990.104 |
| Rdh16f1 | 3.645 | 0.268 | 0 | 163.927 |
| Gm47160 | -1.734 | 0.130 | 0 | 967.490 |
| Ctnnal1 | 1.810 | 0.140 | 0 | 729.556 |

| symbol | log2FC | lfcSE | padj | baseMean |
|---|---|---|---|---|
| Ptprn | 1.244 | 0.096 | 0 | 2858.402 |
| Prok1 | 2.905 | 0.230 | 0 | 223.664 |
| Hk2 | 1.208 | 0.095 | 0 | 15966.140 |
| Klhl29 | 1.230 | 0.100 | 0 | 1763.498 |
| Nin | 1.269 | 0.104 | 0 | 3036.363 |
| Clu | 0.991 | 0.082 | 0 | 11385.931 |
| Gm3716 | 3.462 | 0.291 | 0 | 249.939 |
| Gfap | 1.310 | 0.112 | 0 | 861.405 |
| Glb1l3 | -4.593 | 0.402 | 0 | 84.684 |
| Bst1 | 1.761 | 0.157 | 0 | 393.738 |
| Tmsb10 | -1.494 | 0.133 | 0 | 1720.990 |
| Gm41608 | 2.743 | 0.259 | 0 | 229.377 |
| Agtpbp1 | 0.952 | 0.089 | 0 | 5351.841 |

#### Kcnv2 in the fitted model

▶ Code

Kcnv2 in the retained-sample model

| symbol | baseMean | log2FC | lfcSE | pvalue | padj |
|---|---|---|---|---|---|
| Kcnv2 | 9599.614 | -0.2756 | 0.0697 | 0 | 5e-04 |

### Volcano plot

▶ Code

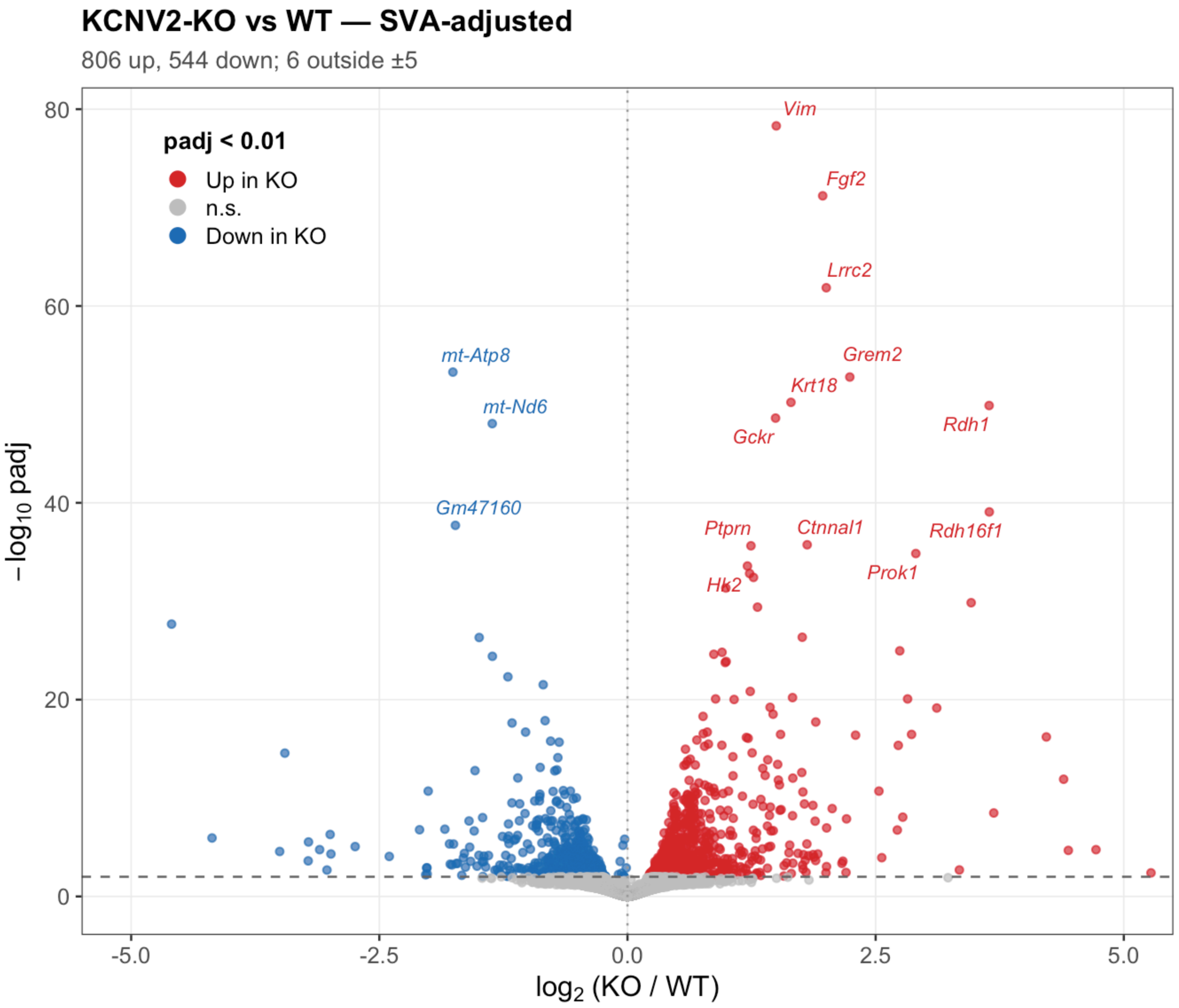


#### Diagnostics

▶ Code

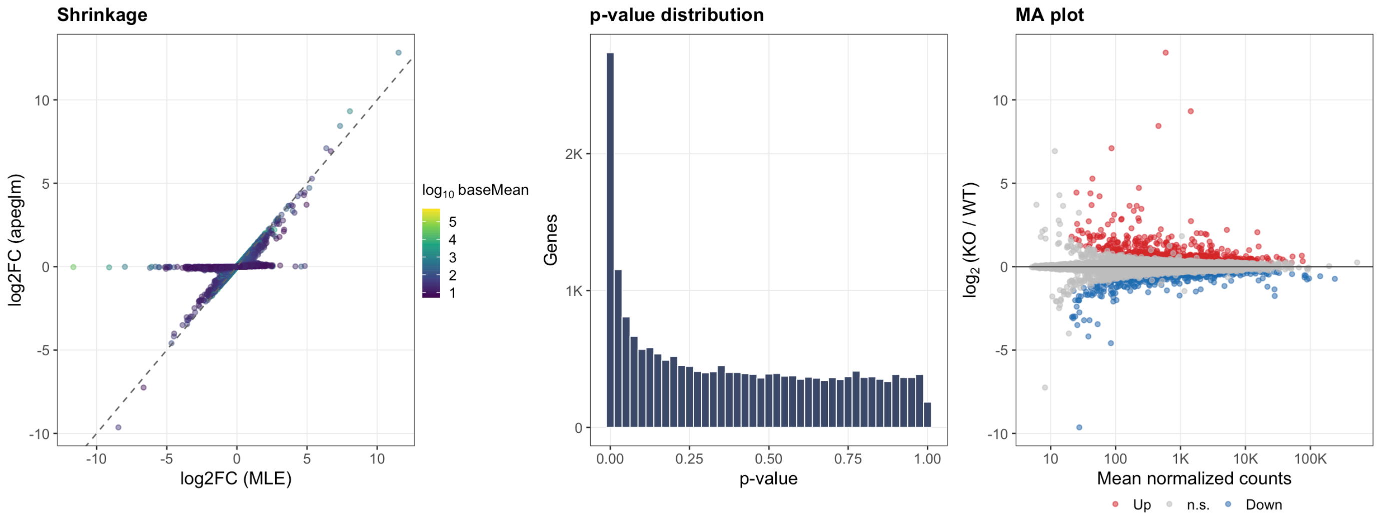


▶ Code

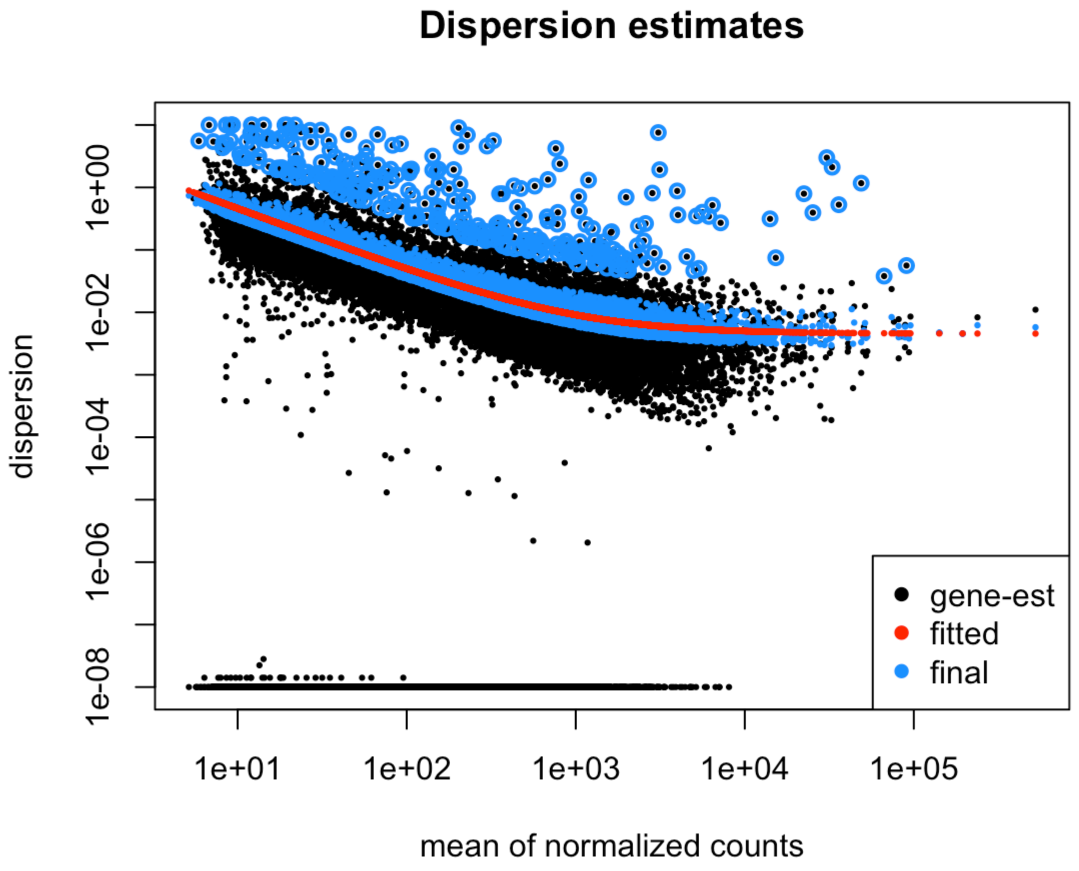


#### Heatmap of top differential genes

▶ Code

▶ Code

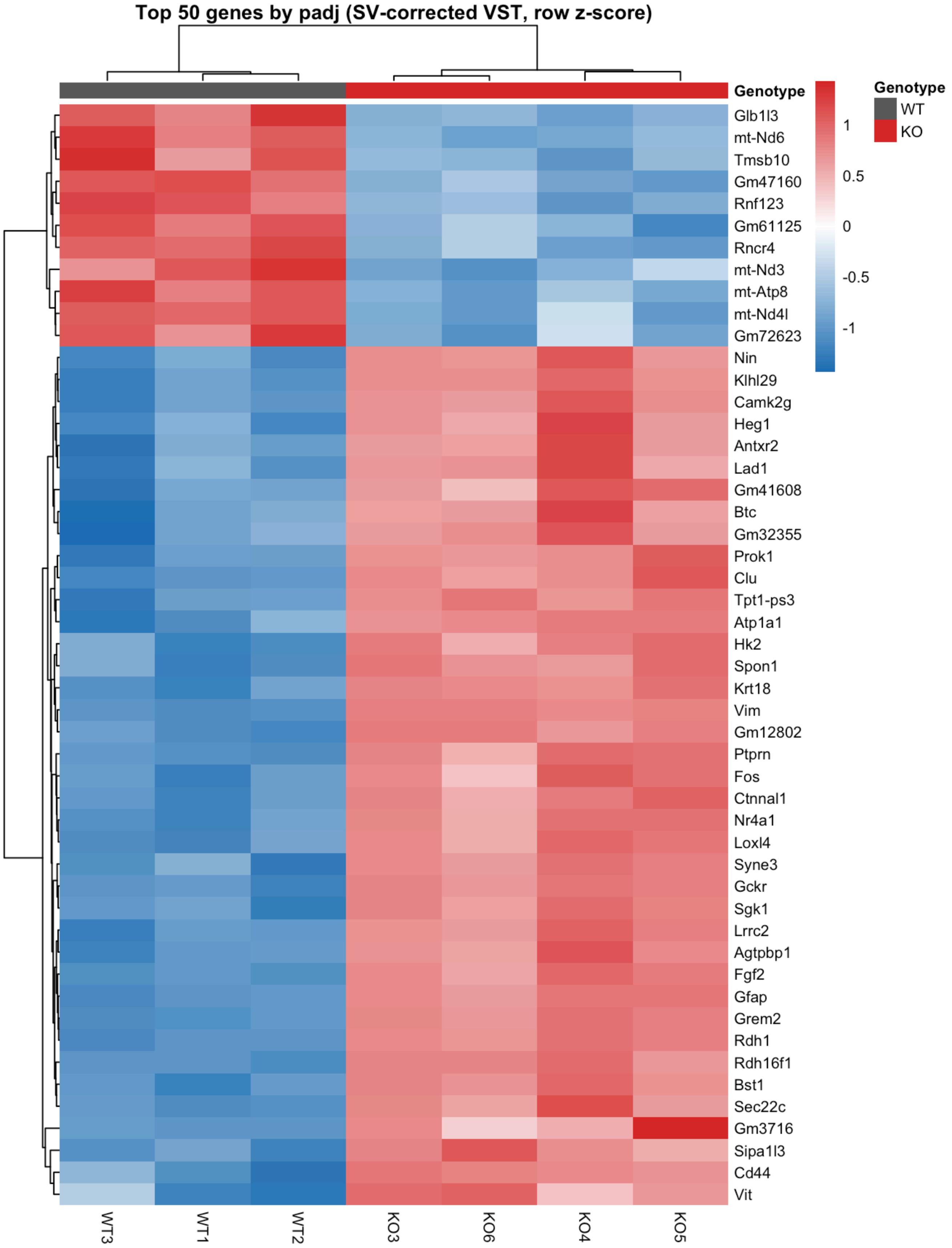


#### Osmosensitive gene panel

The gene set from Panel B of the reference figure. Bars are log2FC with posterior SD; fill is the observed direction at `padj < 0.05`, stars are `***` padj < 0.001, `**` < 0.01, `*` < 0.05.

▶ Code

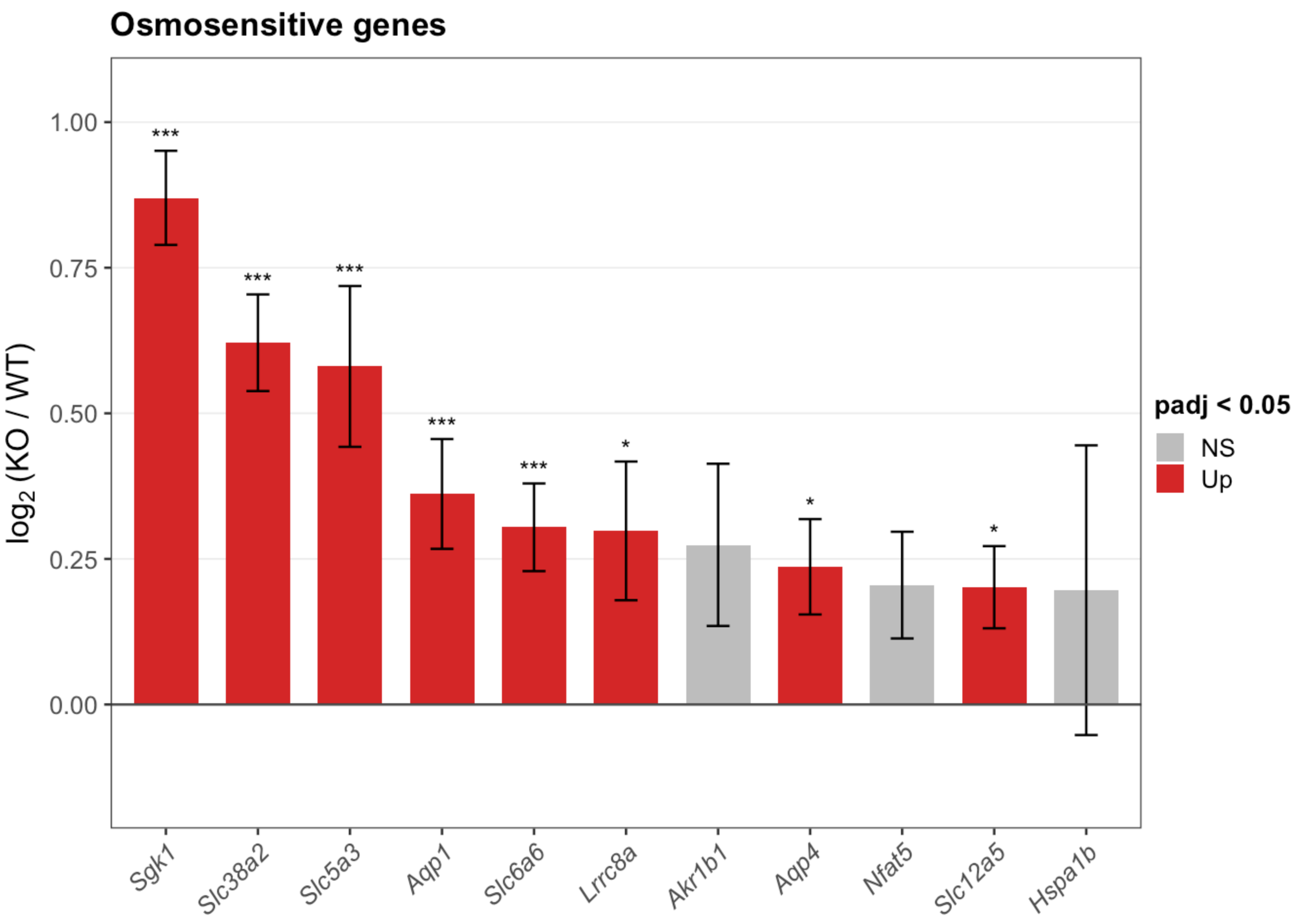


#### Müller glia gliosis signature

Reactive gliosis is a **bidirectional** shift: Müller glia up-regulate a stress and intermediate-filament program while down-regulating the homeostatic machinery that defines the resting cell. Showing only the induced half overstates the case, so both tiers are plotted together — the signature is convincing when the left panel rises *and* the right panel falls.

Bars are log2FC with posterior SD; stars are `***` padj < 0.001, `**` < 0.01, `*` < 0.05. Fill encodes the actual result (red up, blue down, grey n.s.), which is deliberately independent of the tier a gene was assigned to — a homeostatic marker coming out red is visible immediately.

▶ Code

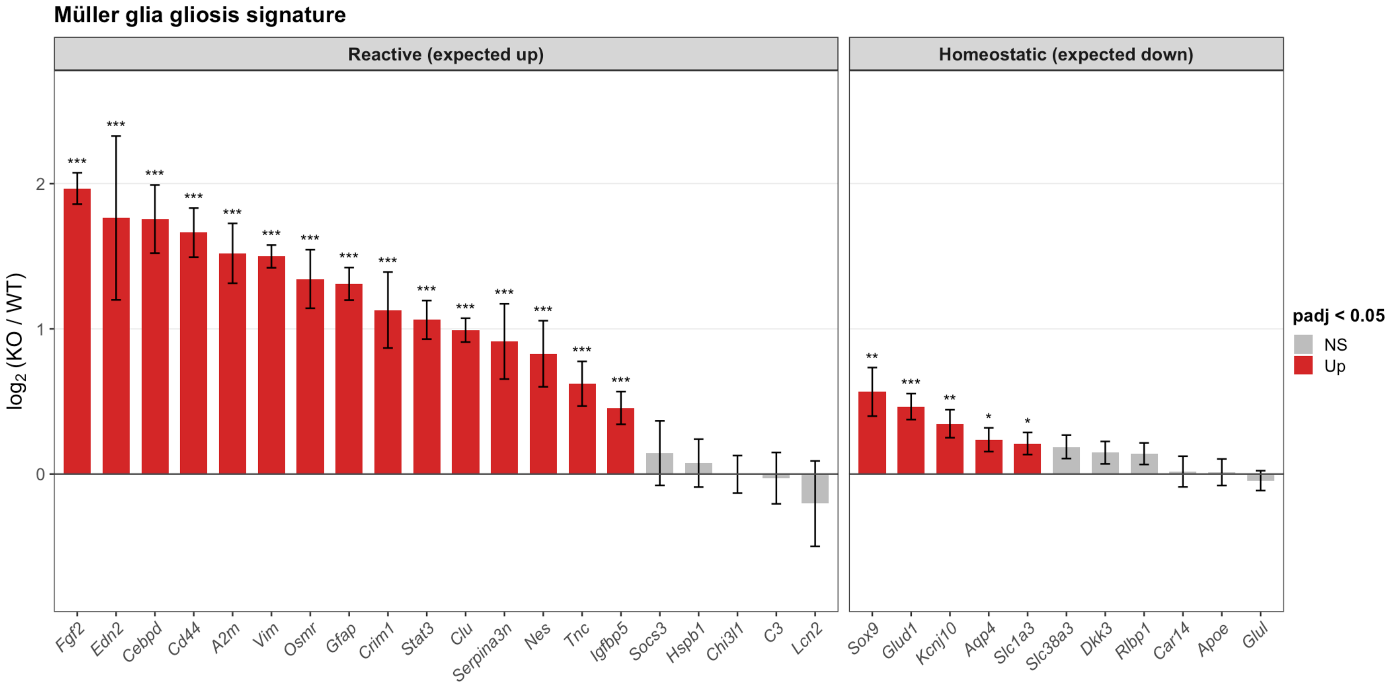


▶ Code

Gliosis tiers vs. transcriptome background (Wilcoxon)

| tier | n_genes | median_lfc | n_sig_up | n_sig_down | wilcox_p |
|---|---|---|---|---|---|
| Reactive (expected up) | 20 | 1.026 | 15 | 0 | 0.000000 |
| Homeostatic (expected down) | 11 | 0.188 | 5 | 0 | 0.000454 |

#### Retinal cell-type marker panels

Whole-retina RNA-seq cannot distinguish a change in transcription within cells from a change in the cellular composition of the tissue. Coordinated movement of an entire marker panel argues for composition.

▶ Code

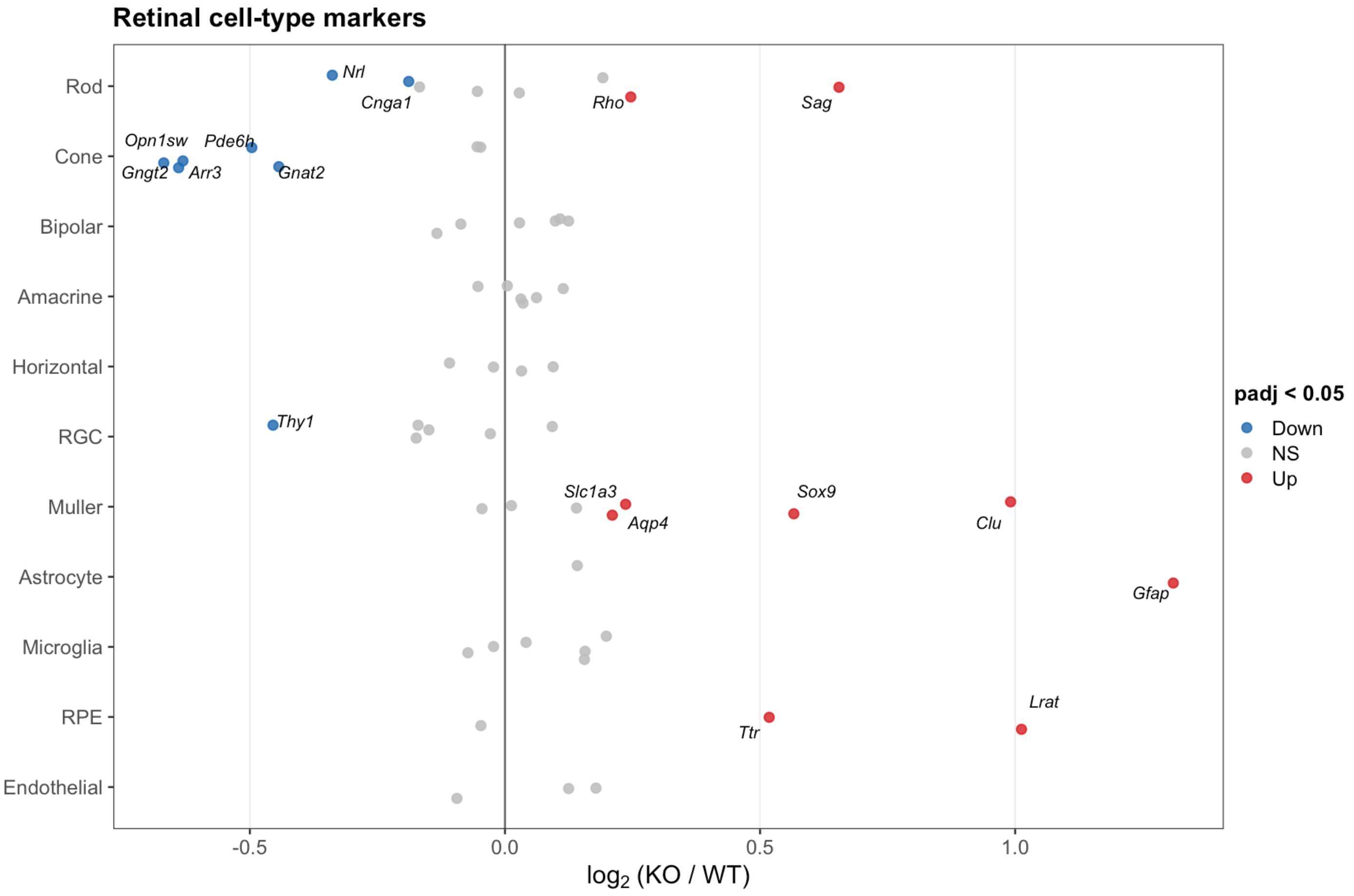


▶ Code

Cell-type marker summary

| cell_type | n_genes | median_lfc | n_up | n_down |
|---|---|---|---|---|
| Astrocyte | 2 | 0.726 | 1 | 0 |
| RPE | 3 | 0.518 | 2 | 0 |
| Cone | 7 | -0.496 | 0 | 5 |
| Muller | 7 | 0.210 | 4 | 0 |
| RGC | 6 | -0.160 | 0 | 1 |
| Endothelial | 3 | 0.125 | 0 | 0 |
| Microglia | 6 | 0.098 | 0 | 0 |
| Bipolar | 6 | 0.064 | 0 | 0 |
| Amacrine | 6 | 0.033 | 0 | 0 |
| Rod | 8 | -0.013 | 2 | 2 |
| Horizontal | 4 | 0.005 | 0 | 0 |

#### Gene set enrichment (GSEA)

Genes ranked by log2FC. GSEA uses the whole ranked list and needs no significance cutoff, which makes it sensitive to coordinated shifts that never reach per-gene significance.

▶ Code

▶ Code

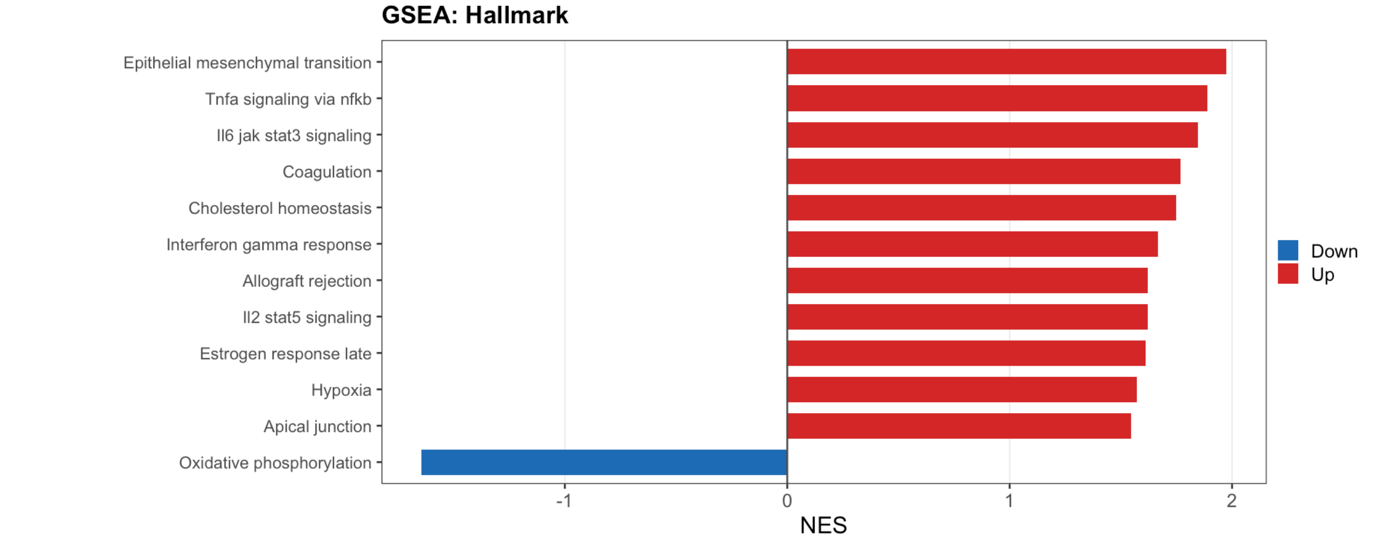


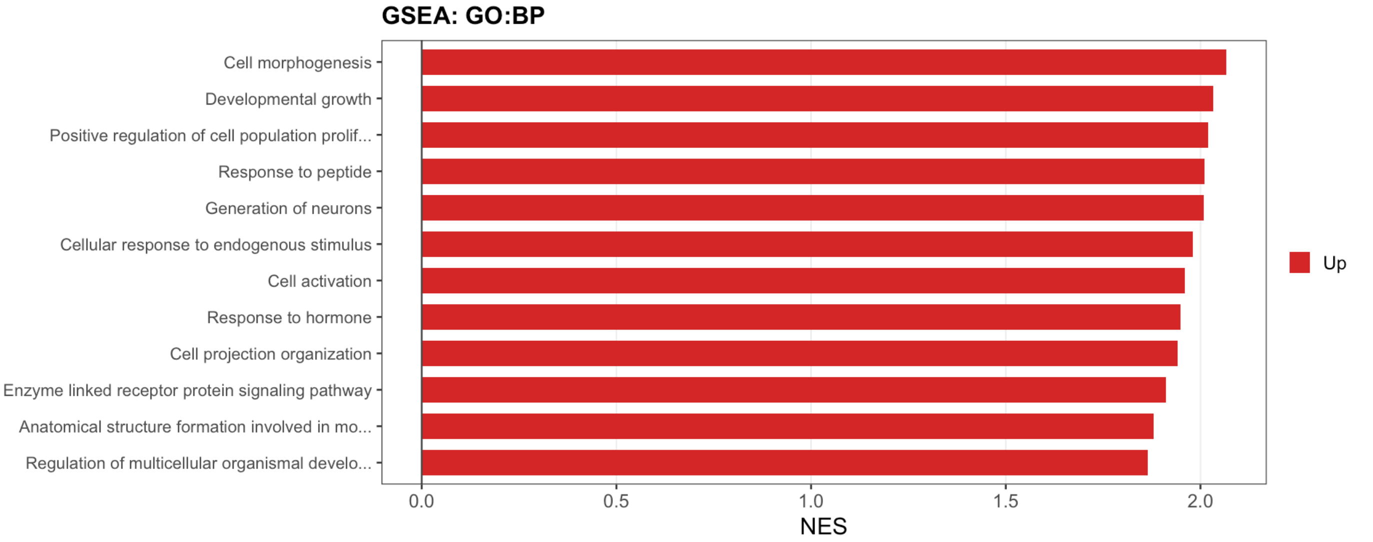


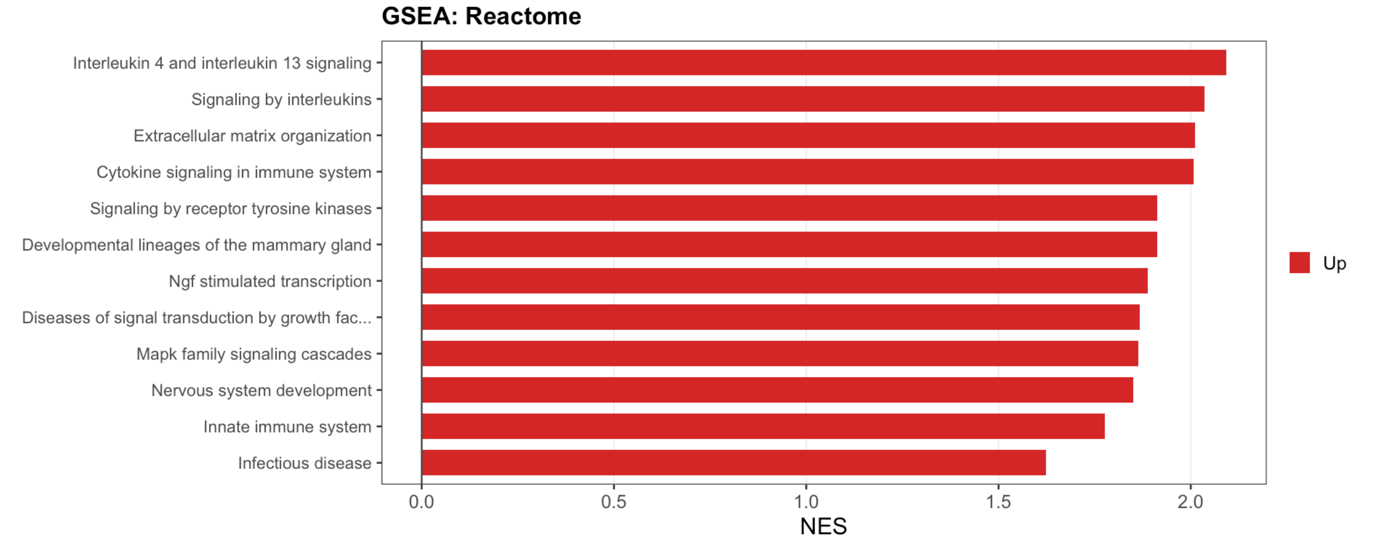

### Pathway over-representation analysis

GSEA and over-representation answer different questions. GSEA asks whether a gene set drifts systematically across the whole ranked list; ORA asks whether the `padj < 0.01` set is enriched for a pathway relative to the tested background. A result supported by both is considerably more secure than one found by either alone.

Four databases are queried, up- and down-regulated genes separately. KEGG and WikiPathways fetch over the network at render time, so each call is wrapped — a failure logs a message and drops that database rather than killing the render.

▶ Code

```
Universe: 14592 genes
```

▶ Code

```
Up: 756  Down: 470
```

▶ Code

▶ Code

Significantly enriched terms per database (padj < 0.05)

| database | down | up |
|---|---:|---:|
| GO:BP | 10 | 199 |
| GO:CC | 8 | 44 |
| GO:MF | 4 | 26 |
| Reactome | 16 | 39 |

▶ Code

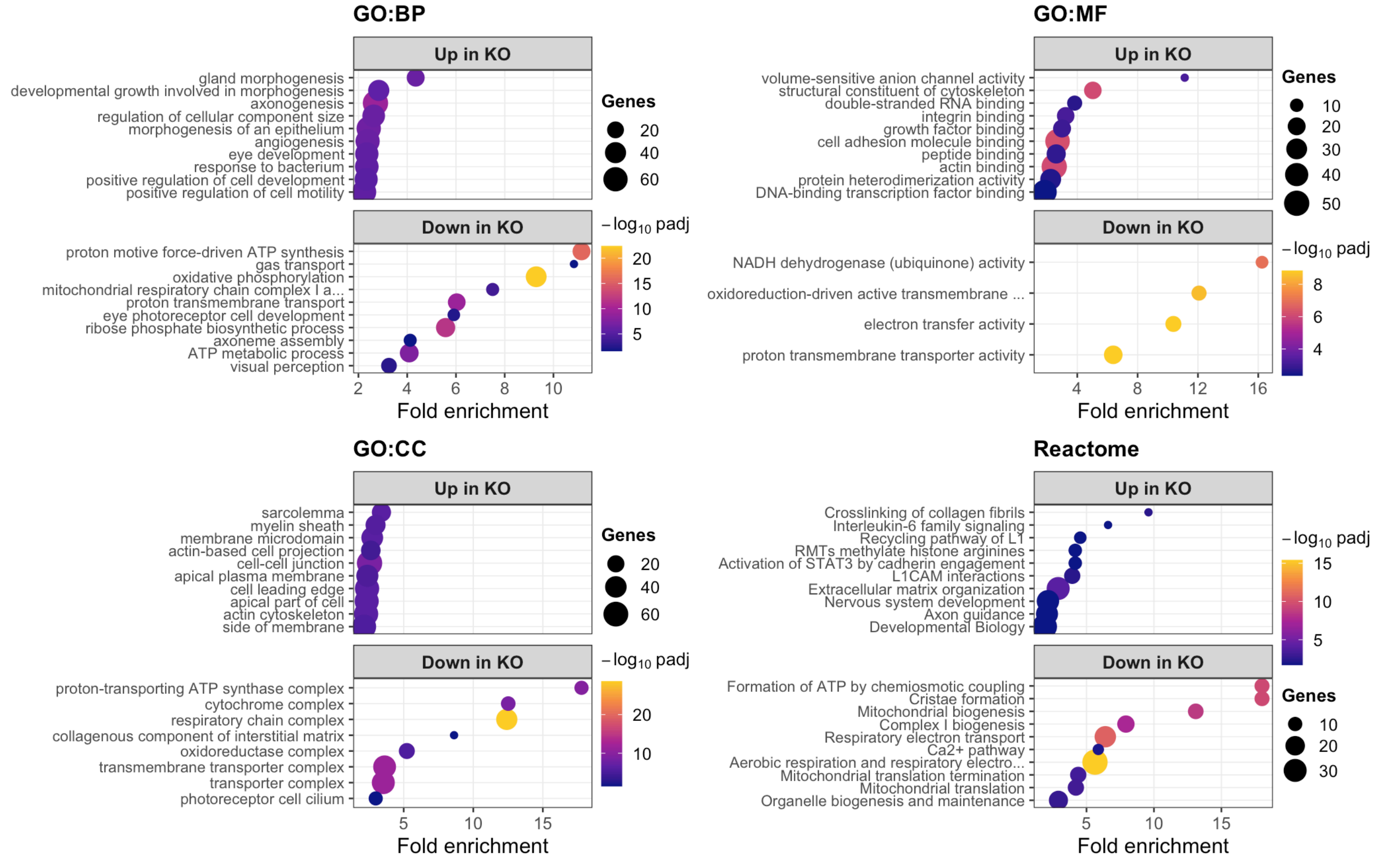


▶ Code

Top 3 enriched terms per database and direction

| database | direction | Description | fold_enrichment | Count | p.adjust |
|---|---|---|---|---|---|
| GO:BP | down | oxidative phosphorylation | 9.29 | 38 | 0.000000 |
| GO:BP | down | proton motive force-driven ATP synthesis | 11.15 | 24 | 0.000000 |
| GO:BP | down | ribose phosphate biosynthetic process | 5.58 | 30 | 0.000000 |
| GO:BP | up | axonogenesis | 2.70 | 67 | 0.000000 |
| GO:BP | up | morphogenesis of an epithelium | 2.42 | 60 | 0.000000 |
| GO:BP | up | angiogenesis | 2.38 | 59 | 0.000001 |
| GO:CC | down | respiratory chain complex | 12.41 | 38 | 0.000000 |
| GO:CC | down | transmembrane transporter complex | 3.63 | 47 | 0.000000 |
| GO:CC | down | transporter complex | 3.53 | 48 | 0.000000 |
| GO:CC | up | cell-cell junction | 2.54 | 63 | 0.000000 |
| GO:CC | up | cell leading edge | 2.36 | 56 | 0.000000 |
| GO:CC | up | apical part of cell | 2.34 | 55 | 0.000000 |
| GO:MF | down | proton transmembrane transporter activity | 6.38 | 22 | 0.000000 |
| GO:MF | down | electron transfer activity | 10.37 | 15 | 0.000000 |

| database | direction | Description | fold_enrichment | Count | p.adjust |
|---|---|---|---|---|---|
| GO:MF | down | oxidoreduction-driven active transmembrane transporter activity | 12.07 | 13 | 0.000000 |
| GO:MF | up | cell adhesion molecule binding | 2.69 | 46 | 0.000001 |
| GO:MF | up | actin binding | 2.47 | 50 | 0.000001 |
| GO:MF | up | structural constituent of cytoskeleton | 5.03 | 19 | 0.000001 |
| Reactome | down | Aerobic respiration and respiratory electron transport | 5.63 | 38 | 0.000000 |
| Reactome | down | Respiratory electron transport | 6.40 | 25 | 0.000000 |
| Reactome | down | Formation of ATP by chemiosmotic coupling | 18.01 | 11 | 0.000000 |
| Reactome | up | Extracellular matrix organization | 2.90 | 30 | 0.000083 |
| Reactome | up | Crosslinking of collagen fibrils | 9.60 | 6 | 0.004740 |
| Reactome | up | L1CAM interactions | 3.94 | 13 | 0.004740 |

#### Where GSEA and ORA agree

▶ Code

Terms called by both GSEA and ORA, same direction

| term | gsea_dir | NES | gsea_padj | ora_dir | database | fold_enrichment | ora_padj |
|---|---|---|---|---|---|---|---|
| cytokine signaling in immune system | up | 2.008 | 0.000 | up | Reactome | 1.732 | 0.040 |
| cell junction organization | up | 1.916 | 0.000 | up | Reactome | 2.143 | 0.021 |
| positive regulation of cell adhesion | up | 2.072 | 0.000 | up | GO:BP | 2.245 | 0.000 |
| cytokine mediated signaling pathway | up | 2.068 | 0.001 | up | GO:BP | 1.715 | 0.026 |
| immune effector process | up | 1.900 | 0.003 | up | GO:BP | 1.815 | 0.006 |
| response to bacterium | up | 1.925 | 0.004 | up | GO:BP | 2.342 | 0.000 |
| signaling by interleukins | up | 2.035 | 0.004 | up | Reactome | 2.119 | 0.024 |
| signaling by receptor tyrosine kinases | up | 1.913 | 0.005 | up | Reactome | 1.724 | 0.035 |
| regulation of actin filament based process | up | 1.967 | 0.006 | up | GO:BP | 2.088 | 0.000 |
| regulation of supramolecular fiber organization | up | 1.903 | 0.006 | up | GO:BP | 2.112 | 0.000 |
| extracellular matrix organization | up | 2.011 | 0.007 | up | GO:BP | 2.328 | 0.000 |
| actin filament organization | up | 1.957 | 0.007 | up | GO:BP | 2.084 | 0.000 |
| protein localization to nucleus | up | 1.987 | 0.008 | up | GO:BP | 1.685 | 0.034 |
| regulation of cell activation | up | 1.748 | 0.009 | up | GO:BP | 1.649 | 0.021 |
| nervous system development | up | 1.851 | 0.010 | up | Reactome | 2.142 | 0.021 |
| cellular response to lipid | up | 1.781 | 0.010 | up | GO:BP | 1.833 | 0.002 |
| erk1 and erk2 cascade | up | 1.948 | 0.010 | up | GO:BP | 2.661 | 0.000 |
| cell junction assembly | up | 1.883 | 0.010 | up | GO:BP | 1.744 | 0.004 |

| term | gsea_dir | NES | gsea_padj | ora_dir | database | fold_enrichment | ora_padj |
|---|---|---|---|---|---|---|---|
| regulation of developmental growth | up | 1.953 | 0.011 | up | GO:BP | 2.222 | 0.000 |
| response to wounding | up | 1.787 | 0.012 | up | GO:BP | 2.255 | 0.000 |

#### Counts for individual genes

▶ Code

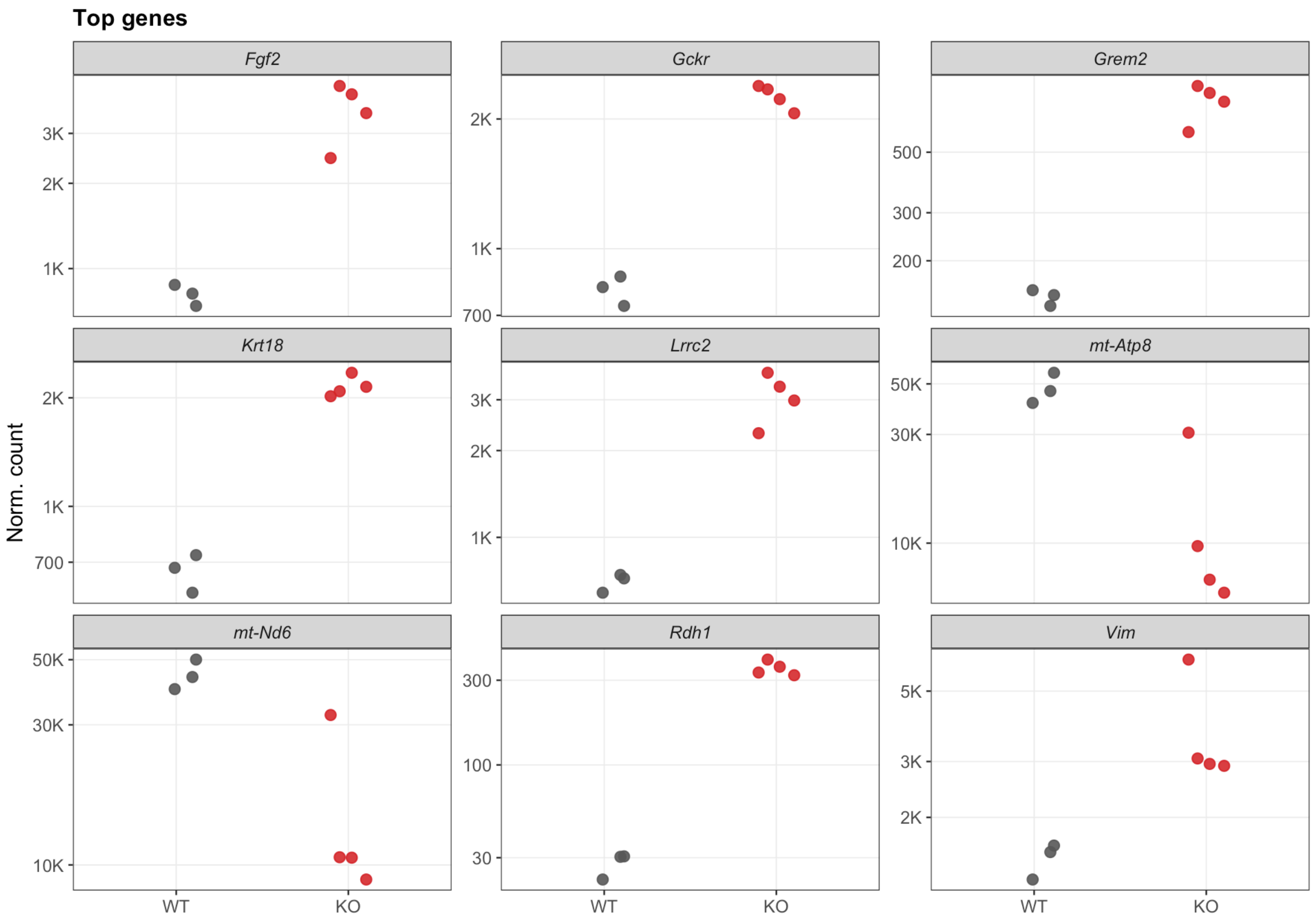


#### Appendix: why SVA

Revision 4 ran every analysis twice, with and without the surrogate variable, to test whether the adjustment was absorbing genotype signal rather than noise. The comparison is condensed here as justification for the modeling choice.

▶ Code

Genes called at padj < 0.01

| model | n_sig |
| --- | --- |
| No SVA | 1315 |
| SVA | 1350 |

▶ Code

Agreement between the two models

| called | n |
| --- | --- |
| Both | 1081 |
| Neither | 18591 |
| No SVA only | 230 |
| SVA only | 231 |

▶ Code

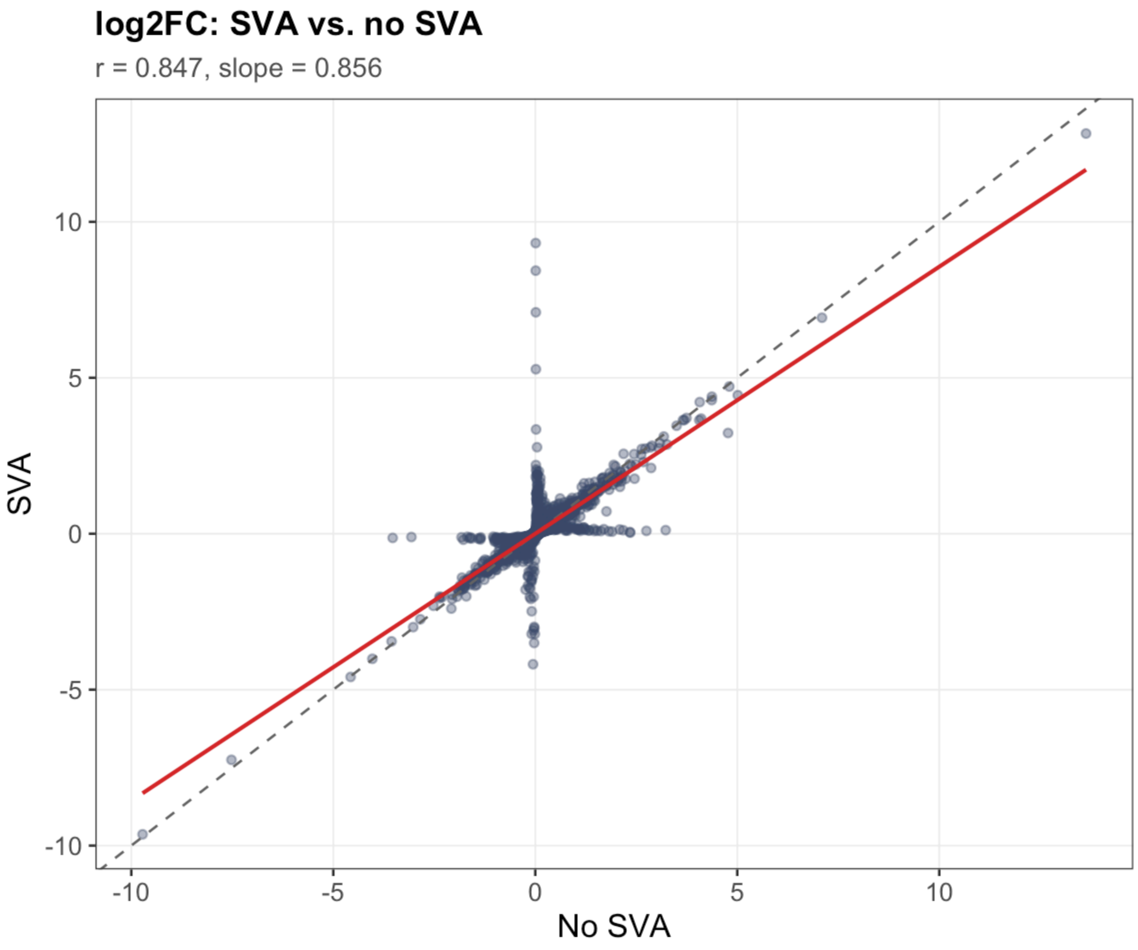


### Figure inventory

▶ Code

```
Figures written to 'figures_pub/'. Square panels are 4 x 4 cm; multi-panel and pathway
figures are wider where a 4 cm axis cannot hold the labels legibly.
```

▶ Code

Publication figures

| file | size_kb |
|---|---|
| appendix_lfc_scatter.pdf | 616.8 |
| compare_called_by_model.pdf | 54.9 |
| compare_gsea_nes.pdf | 115.2 |
| compare_lfc_scatter.pdf | 615.3 |
| compare_padj_scatter.pdf | 350.9 |
| compare_sig_counts.pdf | 15.0 |
| counts_top_genes.pdf | 21.8 |
| counts_top_genes_SVA.pdf | 22.2 |
| counts_top_genes_noSVA.pdf | 22.8 |
| diag_dispersion.pdf | 1563.1 |
| diag_dispersion_SVA.pdf | 1582.7 |
| diag_dispersion_noSVA.pdf | 1588.6 |
| diag_ma_plot.pdf | 627.6 |
| diag_ma_plot_SVA.pdf | 645.0 |
| diag_ma_plot_noSVA.pdf | 648.2 |
| diag_pvalue_hist.pdf | 14.2 |
| diag_pvalue_hist_SVA.pdf | 13.4 |
| diag_pvalue_hist_noSVA.pdf | 13.8 |
| diag_shrinkage.pdf | 655.3 |
| gsea_GOBP.pdf | 15.4 |
| gsea_GOBP_SVA.pdf | 19.9 |
| gsea_GOBP_noSVA.pdf | 16.2 |
| gsea_Hallmark.pdf | 16.1 |
| gsea_Hallmark_SVA.pdf | 15.6 |
| gsea_Hallmark_noSVA.pdf | 15.8 |
| gsea_Reactome.pdf | 17.0 |
| gsea_Reactome_SVA.pdf | 16.7 |
| gsea_Reactome_noSVA.pdf | 17.2 |
| heatmap_top25.pdf | 6.6 |

| file | size_kb |
|---|---|
| heatmap_top25_SVA.pdf | 6.8 |
| heatmap_top25_noSVA.pdf | 6.8 |
| ko_validation_kcnv2.pdf | 14.6 |
| ora_GOBP.pdf | 17.7 |
| ora_GOBP_SVA.pdf | 27.2 |
| ora_GOBP_noSVA.pdf | 26.9 |
| ora_GOCC.pdf | 16.2 |
| ora_GOMF.pdf | 17.7 |
| ora_Reactome.pdf | 23.1 |
| panel_gliosis.pdf | 30.1 |
| panel_osmosensitive.pdf | 23.9 |
| panel_osmosensitive_SVA.pdf | 22.1 |
| panel_osmosensitive_noSVA.pdf | 22.6 |
| panel_retina_markers.pdf | 19.7 |
| panel_retina_markers_SVA.pdf | 17.5 |
| panel_retina_markers_noSVA.pdf | 17.9 |
| qc_cooks_distance.pdf | 164.5 |
| qc_genes_detected.pdf | 14.1 |
| qc_mapped_reads.pdf | 13.8 |
| qc_mapping_rate.pdf | 14.7 |
| qc_pca_retained.pdf | 14.4 |
| qc_pca_sv_corrected.pdf | 13.7 |
| qc_pca_uncorrected.pdf | 14.2 |
| qc_sample_distances.pdf | 0.0 |
| qc_sv_loadings.pdf | 13.7 |
| shrinkage_mle_vs_apeglm_SVA.pdf | 698.8 |
| shrinkage_mle_vs_apeglm_noSVA.pdf | 693.8 |
| volcano.pdf | 354.6 |
| volcano_SVA.pdf | 375.2 |
| volcano_SVA_nolabels.pdf | 367.6 |
| volcano_noSVA.pdf | 351.3 |
| volcano_noSVA_nolabels.pdf | 343.7 |
| volcano_nolabels.pdf | 347.1 |

### Session info

▶ Code

```
R version 4.6.1 (2026-06-24)
Platform: aarch64-apple-darwin23
Running under: macOS Tahoe 26.5.2

Matrix products: default
BLAS:   /Library/Frameworks/R.framework/Versions/4.6/Resources/lib/libRblas.0.dylib
LAPACK: /Library/Frameworks/R.framework/Versions/4.6/Resources/lib/libRlapack.dylib;
LAPACK version 3.12.1

locale:
[1] en_US.UTF-8/en_US.UTF-8/en_US.UTF-8/C/en_US.UTF-8/en_US.UTF-8

time zone: America/Chicago
tzcode source: internal

attached base packages:
[1] stats4    stats     graphics  grDevices utils     datasets  methods
[8] base

other attached packages:
 [1] lubridate_1.9.5             forcats_1.0.1
 [3] stringr_1.6.0               dplyr_1.2.1
 [5] purrr_1.2.2                 readr_2.2.0
 [7] tidyr_1.3.2                 tibble_3.3.1
 [9] tidyverse_2.0.0             rjson_0.2.23
[11] scales_1.4.0                openxlsx_4.2.8.1
[13] msigdbr_26.1.0              patchwork_1.3.2
[15] ggpubr_1.0.0                cowplot_1.2.0
[17] ggrepel_0.9.8               ggplot2_4.0.3
[19] viridis_0.6.5               viridisLite_0.4.3
[21] RColorBrewer_1.1-3          pheatmap_1.0.13
[23] vsn_3.80.0                  fgsea_1.38.0
[25] ReactomePA_1.56.0           DOSE_4.6.0
[27] enrichplot_1.32.0           clusterProfiler_4.20.0
[29] biomaRt_2.68.0              org.Mm.eg.db_3.23.0
[31] txdbmaker_1.7.3             GenomicFeatures_1.64.0
[33] AnnotationDbi_1.74.0        limma_3.68.4
[35] sva_3.60.0                  BiocParallel_1.46.0
[37] genefilter_1.94.0           mgcv_1.9-4
[39] nlme_3.1-170                apeglm_1.34.0
[41] DESeq2_1.52.0               SummarizedExperiment_1.42.0
[43] Biobase_2.72.0              MatrixGenerics_1.24.0
[45] matrixStats_1.5.0           GenomicRanges_1.64.0
[47] Seqinfo_1.2.0               IRanges_2.46.0
[49] S4Vectors_0.50.1            BiocGenerics_0.58.1
[51] generics_0.1.4              tximport_1.40.0
```

```
loaded via a namespace (and not attached):
  [1] fs_2.1.0                 bitops_1.1-0             httr_1.4.8
  [4] numDeriv_2016.8-1.1      tools_4.6.1              backports_1.5.1
  [7] R6_2.6.1                 lazyeval_0.2.3           withr_3.0.3
 [10] graphite_1.58.0          prettyunits_1.2.0        gridExtra_2.3.1
 [13] preprocessCore_1.74.0    cli_3.6.6                scatterpie_0.2.6
 [16] labeling_0.4.3           mvtnorm_1.4-2            S7_0.2.2
 [19] Rsamtools_2.28.0         systemfonts_1.3.2        yulab.utils_0.2.4
 [22] gson_0.2.0               dichromat_2.0-1          bbmle_1.0.25.1
 [25] rstudioapi_0.19.0        RSQLite_3.53.3           gridGraphics_0.5-1
 [28] BiocIO_1.22.0            vroom_1.7.1              zip_3.0.1
 [31] car_3.1-5                GO.db_3.23.1             Matrix_1.7-6
 [34] abind_1.4-8              lifecycle_1.0.5          yaml_2.3.12
 [37] edgeR_4.10.1             carData_3.0-6            qvalue_2.44.0
 [40] SparseArray_1.12.2       BiocFileCache_3.2.0      grid_4.6.1
 [43] blob_1.3.0               crayon_1.5.3             bdsmatrix_1.3-7
 [46] ggtangle_0.1.2           lattice_0.22-9           cigarillo_1.2.1
 [49] annotate_1.90.0          KEGGREST_1.52.0          pillar_1.11.1
 [52] knitr_1.51               codetools_0.2-20         fastmatch_1.1-8
 [55] glue_1.8.1               ggiraph_0.9.6            ggfun_0.2.1
 [58] fontLiberation_0.1.0     data.table_1.18.4        vctrs_0.7.3
 [61] png_0.1-9                treeio_1.36.1            gtable_0.3.6
 [64] assertthat_0.2.1         emdbook_1.3.14           cachem_1.1.0
 [67] xfun_0.60                S4Arrays_1.12.0          tidygraph_1.3.1
 [70] coda_0.19-4.1            survival_3.8-9           aisdk_1.4.12
 [73] statmod_1.5.2            ggtree_4.2.0             bit64_4.8.2
 [76] fontquiver_0.2.1         progress_1.2.3           filelock_1.0.3
 [79] GenomeInfoDb_1.48.0      affyio_1.82.0            otel_0.2.0
 [82] DBI_1.3.0                tidyselect_1.2.1         processx_3.9.0
 [85] bit_4.6.0                compiler_4.6.1           curl_7.1.0
 [88] httr2_1.3.0              graph_1.90.0             xml2_1.6.0
 [91] fontBitstreamVera_0.1.1  DelayedArray_0.38.2      rtracklayer_1.72.0
 [94] affy_1.90.0              callr_3.8.0              rappdirs_0.3.4
 [97] digest_0.6.39            rmarkdown_2.31           XVector_0.52.0
[100] htmltools_0.5.9          pkgconfig_2.0.3          dbplyr_2.6.0
[103] fastmap_1.2.0            rlang_1.3.0              htmlwidgets_1.6.4
[106] UCSC.utils_1.7.1         farver_2.1.2             jsonlite_2.0.0
[109] GOSemSim_2.38.3          RCurl_1.98-1.19          magrittr_2.0.5
[112] Formula_1.2-5            ggplotify_0.1.3          Rcpp_1.1.2
[115] babelgene_22.9           ape_5.8-1                ggnewscale_0.5.2
[118] gdtools_0.5.1            stringi_1.8.7            ggraph_2.2.2
[121] MASS_7.3-66              plyr_1.8.9               parallel_4.6.1
[124] Biostrings_2.80.1        graphlayouts_1.2.5       splines_4.6.1
[127] hms_1.1.4                locfit_1.5-9.12          igraph_2.3.3
[130] ggsignif_0.6.4           enrichit_0.2.0           reshape2_1.4.5
[133] XML_3.99-0.23            evaluate_1.0.5           BiocManager_1.30.27
[136] tzdb_0.5.0               tweenr_2.0.3             polyclip_1.10-7
[139] ggforce_0.5.0            broom_1.0.13             xtable_1.8-8
[142] restfulr_0.0.17          reactome.db_1.96.0       tidytree_0.4.8
[145] tidydr_0.0.6             rstatix_1.1.0            aplot_0.3.1
```

```
[148] memoise_2.0.1           GenomicAlignments_1.48.0 cluster_2.1.8.3
[151] timechange_0.4.0
```
