## Supplemental File S1 for "Biomarkers indicate altered outer retinal fluid homeostasis in the absence of Kv2.1/Kv8.2 channels"

**Statistical outputs for OCT data (Tables S1A-S4C)**

#### Method

We used a linear mixed model with fixed effects of genotype, condition, side, and all interactions. The model included a random intercept for mouse nested within genotype. The test of interest was the interaction between genotype and condition. We used the model to estimate means for all conditions.

#### 1. ELM-RPE

**Table S1A.** ELM-RPE: Type III ANOVA results (genotype × condition)

| Term | Sum Sq | Mean Sq | NumDF | DenDF | F value | Pr(>F) |
|---|---|---|---|---|---|---|
| genotype | 51.11 | 51.11 | 1 | 15 | 71.40 | 4.36e-07 |
| condition | 56.28 | 56.28 | 1 | 15 | 78.62 | 2.37e-07 |
| genotype:condition | 11.87 | 11.87 | 1 | 15 | 16.58 | 1.00e-03 |

- The main effect of genotype on ELM-RPE thickness was highly significant ($p < 0.0001$).
- There were also significant effects of condition ($p < 0.0001$), the genotype x condition interaction ($p = 0.0010$), side ($p < 0.0001$) and the genotype x side interaction ($p = 0.007$). Therefore, data were analyzed separately by condition and side.

**Table S1B.** ELM-RPE: Type III ANOVA results (genotype × side)

| Term | Sum Sq | Mean Sq | NumDF | DenDF | F value | Pr(>F) |
|---|---|---|---|---|---|---|
| genotype | 25.33 | 25.33 | 1 | 15 | 71.40 | 4.36e-07 |
| side | 22.61 | 22.61 | 1 | 15 | 63.74 | 8.83e-07 |
| genotype:side | 3.40 | 3.40 | 1 | 15 | 9.58 | 7.38e-03 |

**Table S1C.** ELM–RPE: Estimated marginal means.

| Condition | Side | Genotype | Mean | SE | lower CI | upper CI |
|---|---|---|---|---|---|---|
| Dark | Inferior | Het | 52.68 | 0.51 | 51.65 | 53.71 |
| Dark | Inferior | KO | 49.21 | 0.37 | 48.44 | 49.97 |
| Light | Inferior | Het | 56.33 | 0.51 | 55.30 | 57.37 |
| Light | Inferior | KO | 50.11 | 0.37 | 49.35 | 50.87 |
| Dark | Superior | Het | 53.45 | 0.51 | 52.42 | 54.48 |
| Dark | Superior | KO | 51.02 | 0.37 | 50.26 | 51.78 |
| Light | Superior | Het | 57.66 | 0.51 | 56.63 | 58.69 |
| Light | Superior | KO | 53.03 | 0.37 | 52.27 | 53.79 |

**Table S1D.** ELM–RPE: Het–KO contrasts by condition within side.

| Condition | Side | Estimate | lower CI | upper CI | Holm-adj. p |
|---|---|---|---|---|---|
| Dark | Inferior | 3.48 | 2.20 | 4.76 | 3.42e-06 |
| Light | Inferior | 6.23 | 4.95 | 7.51 | 1.26e-11 |
| Dark | Superior | 2.43 | 1.15 | 3.71 | 4.80e-04 |
| Light | Superior | 4.63 | 3.35 | 5.91 | 1.45e-08 |

**Table S1E.** ELM-RPE: Light–Dark contrasts by genotype within side.

| Genotype | Side | Estimate | lower CI | upper CI | Holm-adj. p |
|---|---|---|---|---|---|
| Het | Inferior | 3.65 | 2.62 | 4.68 | 6.56e-09 |
| KO | Inferior | 0.90 | 0.14 | 1.67 | 2.11e-02 |
| Het | Superior | 4.21 | 3.17 | 5.24 | 1.68e-10 |
| KO | Superior | 2.01 | 1.25 | 2.77 | 3.26e-06 |

### 2. $\Delta$ ELM-RPE

- The main effect of genotype on ΔELM-RPE thickness was highly significant ($p < 0.0001$).
- There were also significant effects of side ($p < 0.0001$) and the genotype x side interaction ($p = 0.0074$). Therefore, data were analyzed separately by side.

**Table S2A.** Delta ELM-RPE: Type III ANOVA Results (genotype × side).

| Term | Sum Sq | Mean Sq | NumDF | DenDF | F value | Pr(>F) |
|---|---|---|---|---|---|---|
| genotype | 25.33 | 25.33 | 1 | 15 | 71.40 | 4.36e-07 |
| side | 22.61 | 22.61 | 1 | 15 | 63.74 | 8.83e-07 |
| genotype:side | 3.40 | 3.40 | 1 | 15 | 9.58 | 7.38e-03 |

**Table S2B.** Delta ELM-RPE: Estimated marginal means.

| Side | Genotype | Mean | SE | lower CI | upper CI |
|---|---|---|---|---|---|
| Inferior | Het | 3.65 | 0.52 | 2.55 | 4.75 |
| Inferior | KO | 0.90 | 0.39 | 0.09 | 1.71 |
| Superior | Het | 4.21 | 0.52 | 3.11 | 5.30 |
| Superior | KO | 2.01 | 0.39 | 1.20 | 2.82 |

**Table S2C.** Delta ELM-RPE: Het–KO contrasts within side.

| Side | Estimate | lower CI | upper CI | Holm-adj. p |
|---|---|---|---|---|
| Inferior | 2.75 | 1.38 | 4.11 | 4.49e-04 |
| Superior | 2.20 | 0.84 | 3.56 | 3.12e-03 |

### 3. ONL

**Table S3A.** ONL: Type III ANOVA results (genotype × condition)

| Term | Sum Sq | Mean Sq | NumDF | DenDF | F value | Pr(>F) |
|---|---|---|---|---|---|---|
| genotype | 116.21 | 116.21 | 1 | 15 | 129.64 | 8.83e-09 |
| condition | 0.29 | 0.29 | 1 | 15 | 0.32 | 5.77e-01 |
| genotype:condition | 7.06 | 7.06 | 1 | 15 | 7.88 | 1.33e-02 |

**Table S3B.** ONL: Type III ANOVA results (genotype × side)

| Term | Sum Sq | Mean Sq | NumDF | DenDF | F value | Pr(>F) |
|---|---|---|---|---|---|---|
| genotype | 80.52 | 80.52 | 1 | 15 | 129.64 | 8.83e-09 |
| side | 0.15 | 0.15 | 1 | 15 | 0.24 | 6.32e-01 |
| genotype:side | 13.58 | 13.58 | 1 | 15 | 21.86 | 2.99e-04 |

- The main effect of genotype on ONL thickness was highly significant ($p < 0.0001$).
- The effects of condition ($p = 0.58$) and side ($p = 0.63$) were not significant.
- However, the genotype × condition ($p=0.013$) and genotype × side ($p = 0.0003$) interactions were significant. Therefore, data were analyzed separately by condition and side.

**Table S3C.** ONL: Estimated marginal means.

| Condition | Side | Genotype | Mean | SE | lower CI | upper CI |
|---|---|---|---|---|---|---|
| Dark | Inferior | Het | 63.35 | 0.63 | 62.07 | 64.64 |
| Dark | Inferior | KO | 53.77 | 0.47 | 52.82 | 54.72 |
| Light | Inferior | Het | 62.26 | 0.63 | 60.98 | 63.55 |
| Light | Inferior | KO | 54.74 | 0.47 | 53.79 | 55.69 |
| Dark | Superior | Het | 61.56 | 0.63 | 60.28 | 62.85 |
| Dark | Superior | KO | 54.78 | 0.47 | 53.83 | 55.73 |
| Light | Superior | Het | 61.13 | 0.63 | 59.84 | 62.42 |
| Light | Superior | KO | 56.10 | 0.47 | 55.15 | 57.05 |

**Table S3D.** ONL: Het–KO contrasts by condition within side.

| Condition | Side | Estimate | lower CI | upper CI | Holm-adj. p |
|---|---|---|---|---|---|
| Dark | Inferior | 9.58 | 7.98 | 11.18 | 1.35e-13 |
| Light | Inferior | 7.52 | 5.92 | 9.12 | 6.24e-11 |
| Dark | Superior | 6.79 | 5.19 | 8.39 | 6.96e-10 |
| Light | Superior | 5.03 | 3.43 | 6.63 | 3.33e-07 |

**Table S3E.** ONL: Light–Dark contrasts by genotype within side.

| Genotype | Side | Estimate | lower CI | upper CI | Holm-adj. p |
|---|---|---|---|---|---|
| Het | Inferior | -1.09 | -2.31 | 0.14 | 8.00e-02 |
| KO | Inferior | 0.97 | 0.07 | 1.88 | 3.53e-02 |
| Het | Superior | -0.43 | -1.66 | 0.79 | 4.79e-01 |
| KO | Superior | 1.32 | 0.42 | 2.22 | 5.05e-03 |

### 4. $\triangle$ ONL

**Table S4A.** Delta ONL: Type III ANOVA Results (genotype × side).

| Term | Sum Sq | Mean Sq | NumDF | DenDF | F value | Pr(>F) |
|---|---|---|---|---|---|---|
| genotype | 4.44 | 4.44 | 1 | 15 | 7.88 | 1.33e-02 |
| side | 1.95 | 1.95 | 1 | 15 | 3.46 | 8.25e-02 |
| genotype:side | 0.18 | 0.18 | 1 | 15 | 0.32 | 5.80e-01 |

- The main effect of genotype on ΔONL thickness was significant ($p = 0.013$).
- The main effect of side did not reach statistical significance ($p = 0.083$), and there was no evidence for a genotype × side interaction ($p = 0.58$). Therefore, ΔONL data were analyzed

with genotype as the primary factor; stratification by side was used for consistent visualization purposes.

**Table S4B.** Delta ONL: Estimated marginal means.

| Side | Genotype | Mean | SE | lower CI | upper CI |
|---|---|---|---|---|---|
| Inferior | Het | -1.09 | 0.59 | -2.32 | 0.14 |
| Inferior | KO | 0.97 | 0.43 | 0.07 | 1.88 |
| Superior | Het | -0.43 | 0.59 | -1.66 | 0.80 |
| Superior | KO | 1.32 | 0.43 | 0.41 | 2.23 |

**Table S4C.** Delta ONL: Het–KO contrasts within side.

| Side | Estimate | lower CI | upper CI | Holm-adj. p |
|---|---|---|---|---|
| Inferior | 2.06 | 0.53 | 3.59 | 1.08e-02 |
| Superior | 1.76 | 0.23 | 3.28 | 2.64e-02 |
